# Multimodal single cell sequencing of human diabetic kidney disease implicates chromatin accessibility and genetic background in disease progression

**DOI:** 10.1101/2022.01.28.478204

**Authors:** Parker C. Wilson, Yoshiharu Muto, Haojia Wu, Anil Karihaloo, Sushrut S. Waikar, Benjamin D. Humphreys

## Abstract

Multimodal single cell sequencing is a powerful tool for interrogating cell-specific changes in transcription and chromatin accessibility. We performed single nucleus RNA (snRNA-seq) and assay for transposase accessible chromatin sequencing (snATAC-seq) on human kidney cortex from donors with and without diabetic kidney disease (DKD) to identify altered signaling pathways and transcription factors associated with DKD. Both snRNA-seq and snATAC-seq had an increased proportion of *VCAM1+* injured proximal tubule cells (PT_VCAM1) in DKD samples. PT_VCAM1 has a pro-inflammatory expression signature and transcription factor motif enrichment implicated NFkB signaling. We used stratified linkage disequilibrium score regression to partition heritability of kidney-function-related traits using publicly-available GWAS summary statistics. Cell-specific PT_VCAM1 peaks were enriched for heritability of chronic kidney disease (CKD), suggesting that genetic background may regulate chromatin accessibility and DKD progression. snATAC-seq found cell-specific differentially accessible regions (DAR) throughout the nephron that change accessibility in DKD and these regions were enriched for glucocorticoid receptor (GR) motifs. Changes in chromatin accessibility were associated with decreased expression of insulin receptor, increased gluconeogenesis, and decreased expression of the GR cytosolic chaperone, *FKBP5*, in the diabetic proximal tubule. Cleavage under targets and release using nuclease (CUT&RUN) profiling of GR binding in bulk kidney cortex and an *in vitro* model of the proximal tubule (RPTEC) showed that DAR co-localize with GR binding sites. CRISPRi silencing of GR response elements (GRE) in the *FKBP5* gene body reduced *FKBP5* expression in RPTEC, suggesting that reduced *FKBP5* chromatin accessibility in DKD may alter cellular response to GR. We developed an open-source tool for single cell allele specific analysis (SALSA) to model the effect of genetic background on gene expression. Heterozygous germline single nucleotide variants (SNV) in proximal tubule ATAC peaks were associated with allele-specific chromatin accessibility and differential expression of target genes within cis-coaccessibility networks. Partitioned heritability of proximal tubule ATAC peaks with a predicted allele-specific effect was enriched for eGFR, suggesting that genetic background may modify DKD progression in a cell-specific manner.

## Introduction

Diabetes is the leading cause of end-stage renal disease (ESRD) and a significant contributor to morbidity and mortality (1). An estimated 40% of patients with diabetes develop chronic kidney disease (CKD), which manifests as worsening proteinuria and renal dysfunction (2). Single cell sequencing is a powerful technique that has advanced our understanding of kidney biology (3). Multimodal integration of single nucleus RNA (snRNA-seq) and assay for transpose-accessible chromatin sequencing (snATAC-seq) provides insight into how transcription factors and chromatin-chromatin interactions regulate expression of nearby genes (4). We have performed snRNA-seq and snATAC-seq on kidney cortex from patients with and without type 2 diabetes to identify cell-specific differentially expressed genes and accessible chromatin regions associated with diabetic kidney disease (DKD). We validated key findings from our multimodal analysis with cleavage under targets and release using nuclease (CUT&RUN) and CRISPR interference (CRISPRi) to directly measure transcription factor binding and modify chromatin accessibility of cis-regulatory elements (CRE). Epigenetic regulation of chromatin accessibility may contribute to long-term expression of DKD-related genes in a process termed metabolic memory (5). Our analysis identified cell-specific changes in chromatin accessibility that co-localize with transcription factor binding sites associated with glucose metabolism and corticosteroid signaling in the diabetic nephron.

Type 2 diabetes is characterized by impaired glucose tolerance and insulin resistance, factors which may contribute to the progression of DKD (2). The kidney is an important regulator of blood pressure and glucose levels, which are therapeutic targets that reduce risk of kidney disease progression. In the kidney, the proximal tubule is the primary site for glucose metabolism and plays a dual role in glucose reabsorption and gluconeogenesis (6). Sodium glucose cotransporter 2 inhibitors (SGLT2i) block glucose reabsorption in the proximal tubule and slow progression of DKD (7). SGLT2i may have benefits that extend beyond glycemic control, which has renewed interest in finding additional therapeutic targets in the kidney (7).

Glucocorticoids and mineralocorticoids comprise a class of hormones called corticosteroids produced in the adrenal cortex. Cortisol is the primary endogenous glucocorticoid that binds glucocorticoid receptor (GR). GR is expressed in multiple kidney cell types, including proximal tubule, thick ascending limb, endothelium, and podocytes (3). The principal mineralocorticoid is aldosterone, which regulates sodium reabsorption by binding mineralocorticoid receptor (MR) in the distal nephron. Single cell sequencing of human DKD has shown that corticosteroid-sensitive genes in the thick ascending limb and distal nephron express a transcriptional signature consistent with increased potassium secretion and decreased paracellular calcium and magnesium reabsorption (8). Chronic exposure to endogenous cortisol and long-term treatment with synthetic glucocorticoids has been linked to type 2 diabetes and metabolic syndrome (9,10). GR signaling regulates a wide variety of cellular processes in the kidney, including glucose homeostasis, sodium transport, and inflammation (9). GR activation increases expression of gluconeogenic genes, which drive the synthesis of glucose from non-carbohydrate substrates like lactate, glutamine, and glycerol (11,12). Gluconeogenesis occurs in the kidney and liver and helps to maintain circulating glucose levels. The kidney contributes approximately half of circulating glucose during prolonged fasting and studies have demonstrated that both glucose reabsorption and gluconeogenesis are increased in type 2 diabetes (11).

Glucocorticoid sensitivity is regulated at the tissue level by expression of GR cytosolic chaperones. *FKBP5* encodes a negative regulator of GR signaling called FK506 binding protein 51 (FKBP5) (13). FKBP5 is an Hsp90 co-chaperone that limits GR ligand binding and nuclear translocation to inhibit GR-induced transcriptional responses (13). GR activation stimulates *FKBP5* expression via a negative feedback loop by binding glucocorticoid receptor response elements (GRE) in the *FKBP5* gene body (14). Epigenetic regulation of *FKBP5* is best-characterized as an important modifier of the stress response, but has also been associated with increased cardiometabolic risk in patients with type 2 diabetes (15,16).

Furthermore, *FKBP5* polymorphisms increase susceptibility for obesity-related insulin resistance and hypertriglyceridemia (17,18). These studies suggest that epigenetic regulation and genetic background may modulate GR signaling and cellular stress response in type 2 diabetes (9). Genetic variation contributes to the risk of type 2 diabetes and DKD progression as shown by genome wide association studies (GWAS) (19–21). Many of the variants identified by GWAS are common (MAF > 0.01) and explain a small proportion of heritability of type 2 diabetes and kidney disease (21).

GWAS have been used to investigate a wide variety of kidney-function-related traits, however, one of the difficulties with GWAS is assigning function to risk variants located in non-coding regions (22,23). Recent studies have shown that a significant proportion of GWAS variants for type 1 and type 2 diabetes are located in cell-specific open chromatin regions (24,25). Chromatin accessibility quantitative trait loci (caQTL) are single nucleotide variants (SNV) that correlate with chromatin accessibility in a genomic region (23). A subset of caQTL are also associated with changes in expression of nearby genes (eQTL) (26). The relationship between gene expression and chromatin accessibility can be modeled using allele-specific chromatin accessibility (ASCA) (27–29). Every cell has a maternal and paternal allele that carries a unique genetic fingerprint that may result in altered activity of CRE leading to increased or decreased expression of target genes. For example, the presence of a SNV within a CRE may disrupt a transcription factor binding site on the paternal allele and reduce its enhancer activity. snATAC-seq can directly measure ASCA by quantifying the ratio of ATAC peak fragments that intersect heterozygous germline SNV. snATAC-seq can also predict CRE gene targets with cis-coaccessibility networks (CCAN), which makes it a compelling method for interrogating the effect of genetic variants in open chromatin regions (28). Multimodal single cell datasets can take this approach one step further by integrating snATAC-seq with snRNA-seq to estimate the expression of target genes in single cells as a result of ASCA (4,30). These new methods open the door to novel techniques for gene-enhancer predictions and quantitation of single cell allele-specific effects (30).

In this study, we used snRNA-seq and snATAC-seq to analyze kidney cortex samples obtained from donors with and without DKD. We report cell-specific changes in chromatin accessibility associated with alterations in insulin signaling and glucose metabolism, predominantly in the proximal tubule. Cell-specific changes in chromatin accessibility were corroborated by changes in transcription and enrichment of similar pathways by snRNA-seq. Differentially accessible chromatin regions were enriched for GR motifs and predicted GRE were validated by CUT&RUN in bulk kidney cortex and an *in vitro* model of the proximal tubule. CRISPRi silencing of GRE in the *FKBP5* gene body decreased *FKBP5* expression and implicates decreased chromatin accessibility as a potential mechanism for altering the GR negative feedback loop. Partitioned heritability of cell-specific ATAC peaks and differentially accessible regions in DKD are enriched for heritability of eGFR in the proximal tubule. Single cell allele-specific analysis of SNV in ATAC peaks with SALSA shows that genetic background is associated with changes in expression of target genes and that these ATAC peaks co-localize with the same regions that change accessibility in DKD.

## Results

### Patient Demographics and Clinical Information

A total of 13 kidney cortex samples were obtained from healthy control patients (n=6), and patients with diabetic kidney disease (DKD, n=7). Tissue samples were collected following nephrectomy for renal mass (n=10) or from deceased organ donors (n=3). Patients ranged in age from 50 to 78 years (median=57y) and included seven men and six women (Supplemental Table 1). Patients with type 2 diabetes had elevated hemoglobin A1c (mean=8.2 +/-1.5%). The majority of patients with DKD were on antihypertensive or ACE inhibitor therapy and two patients were on insulin. Two patients with DKD had mild to moderate proteinuria as measured by urine dipstick.

### Renal Histology of Donor Samples

Tissue sections were stained with H&E and examined by a renal pathologist (P.W.) to evaluate histological features of DKD. Control samples did not have significant global glomerulosclerosis (<10%) or interstitial fibrosis and tubular atrophy (<10%). Patients with DKD had predominantly mild (N=3, < 25%) or moderate (N=3, 26-50%) global glomerulosclerosis with a corresponding increase in interstitial fibrosis and tubular atrophy. Mean eGFR of DKD samples (66 +/-25 ml/min/1.73m^2) and control samples (74 +/-15ml/min/1.73m^2) was not statistically different (Students t-test, p=0.49). DKD samples showed nodular mesangial expansion, thickened glomerular basement membranes and afferent arteriolar hyalinosis.

### Single Nucleus ATAC Sequencing in Type 2 Diabetes

The snATAC-seq dataset included six healthy control samples and seven with DKD. snATAC-seq libraries were counted with cellranger-atac (10X Genomics) and aggregated prior to cell-specific peak calling with MACS2 (18,30). We detected 437,311 accessible chromatin regions (‘ATAC peaks’) across all cell types. More abundant cell types had a larger number of ATAC peaks compared to less common cell types, which is likely a function of increased power and sequencing depth (Supplemental Figure 1). The aggregated dataset was analyzed in Signac following batch effect correction with Harmony (30,31). A total of 68,458 cells passed quality control filters and all major cell types in the kidney cortex were represented (Figure 1A). Cell types were identified based on increased chromatin accessibility within gene body and promoter regions of lineage-specific markers (Supplemental Figure 1) and enrichment for cell-specific ATAC peaks (Supplemental Table 2). The most abundant cell type was the proximal convoluted tubule (PCT), which comprised approximately one third of snATAC-seq cells. We previously described an injured population of *VCAM1+* proximal tubule cells (PT_VCAM1) that increase in response to acute kidney injury, aging, and CKD (3). PT_VCAM1 can be distinguished from PCT by expression of *VCAM1* and *HAVCR1* (KIM-1), which is a marker of kidney injury (Supplemental Figure 2). There was a trend towards greater proportion of PT_VCAM1 in DKD samples compared to healthy controls (0.09 vs. 0.25, p = 0.14). DKD samples also had a trend towards greater number of infiltrating leukocytes (mean of 42 vs. 211, p = 0.10), including B cells, T cells, and mononuclear cells.

**Figure 1.**
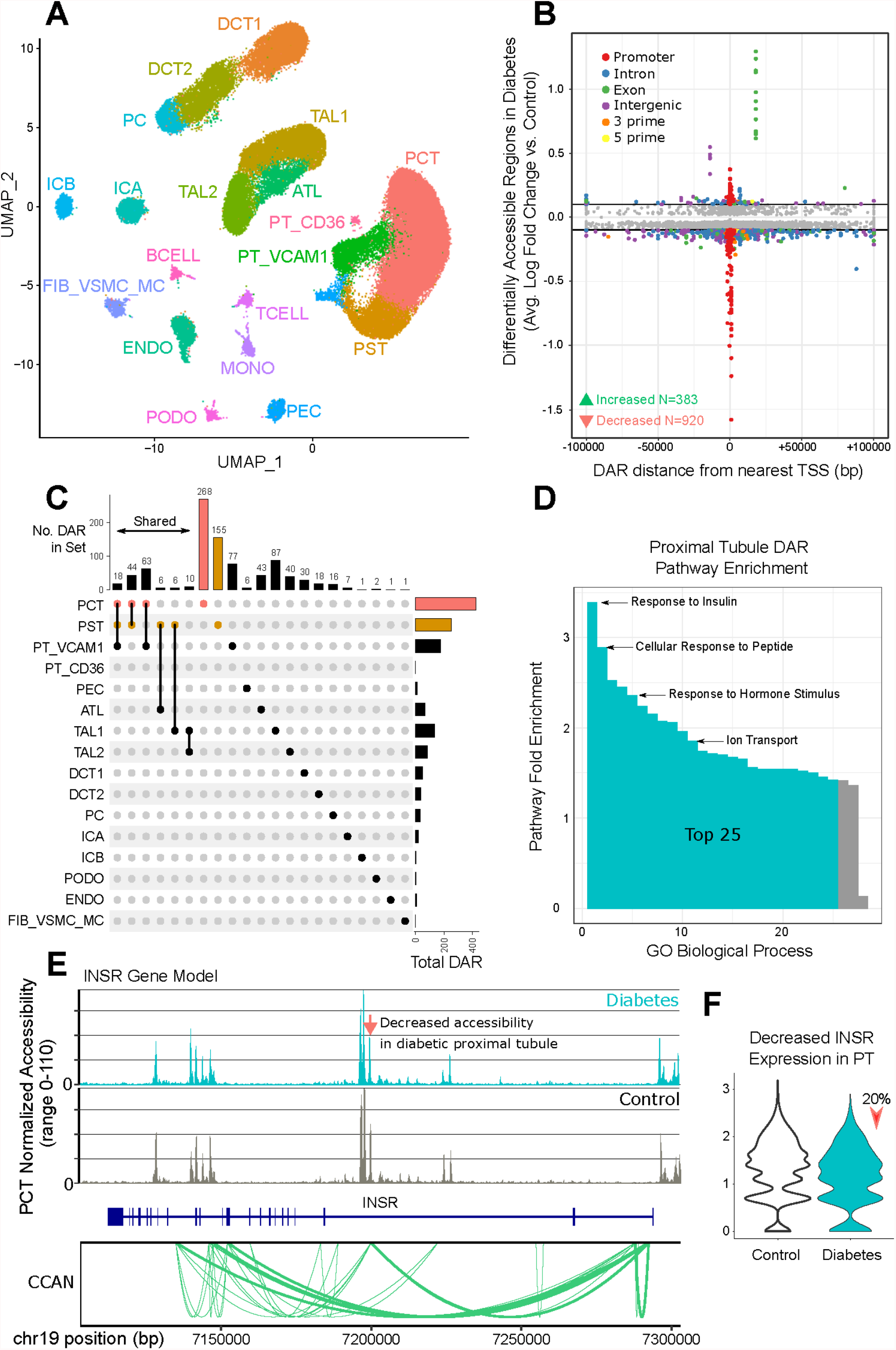
snATAC-seq of human DKD. **A) UMAP embedding of snATAC-seq dataset.** Six healthy control and seven DKD samples were aggregated, preprocessed, and filtered. A total of 68,458 cells are depicted. PCT-proximal convoluted tubule, PST-proximal straight tubule, PT_VCAM1-VCAM1(+) proximal tubule, PT_CD36-CD36(+) proximal tubule cells, PEC-parietal epithelial cells, ATL-ascending thin limb, TAL1-CLDN16(-) thick ascending limb, TAL2-CLDN16(+) thick ascending limb, DCT1-early distal convoluted tubule, DCT2-late distal convoluted tubule, PC-principal cells, ICA-type A intercalated cells, ICB-type B intercalated cells, PODO-podocytes, ENDO-endothelial cells, FIB_VSMC_MC-fibroblasts, vascular smooth muscle cells and mesangial cells, TCELL-T cells, BCELL-B cells, MONO-mononuclear cells. **B) Effect size and location of DAR in DKD**. Healthy control cell types were compared to DKD to identify cell-specific DAR in the snATAC-seq dataset using the Seurat FindMarkers function (Supplemental Table 3). DAR were evaluated with a Bonferroni-adjusted p-value for an FDR < 0.05 and significant DAR that met an absolute log2-fold-change threshold of 0.1 (horizontal bars) were annotated with ChIPSeeker relative to the nearest TSS. **C) DAR in DKD that are cell-specific or shared between cell types**. DAR that were either shared between multiple cell types or unique to a specific cell type are displayed. DAR shared between multiple cell types are limited to groups that share five or more DAR. **D) Proximal tubule DAR pathway enrichment**. Significant cell-specific DAR from PCT and PST were annotated with the nearest protein-coding gene to perform gene ontology enrichment with Panther. Fold-enrichment for all significant GO biological processes is shown and the top 25 are highlighted. **E) Proximal tubule-specific DAR and ATAC peaks in the insulin receptor**. snATAC-seq coverage plots for DKD and control PCT are displayed in relation to the *INSR* gene body. The orange arrow indicates a DAR in intron 2 that shows decreased accessibility in DKD (chr19:7196798-7198626, fold-change=0.92, padj=7.7×10^-13). Green arcs depict the nodes of a cis-coaccessibility network (CCAN) surrounding the *INSR* gene body. **F) Proximal tubule *INSR* expression by snRNA-seq**. Healthy control proximal tubule was compared to DKD in the snRNA-seq dataset to identify differentially expressed genes with the FindMarkers function and visualized as a violin plot. DKD proximal tubule showed reduced *INSR* expression (fold-change=0.78, padj=1.2×10^-27).

### Cell-specific differentially accessible chromatin regions in the diabetic nephron

We compared DKD and control samples to identify 7,347 cell-specific differentially accessible chromatin regions (DAR) that met the adjusted p-value threshold (Supplemental Table 3, Benjamini Hochberg padj < 0.05), including 1,303 that also met an absolute log-fold-change threshold of 0.1 (Figure 1B). The majority of DAR showed decreased accessibility rather than increased accessibility (920 vs. 383) and many were located in a promoter region (Figure 1B). In contrast, a minority of DAR were in intergenic sites (152/1,303, 11%) with a median distance of 50kb from the nearest transcriptional start site (TSS). Nearly half of intergenic DAR (n=69/152, 45%) and one third of intronic DAR (n=64/236, 27%) mapped to a FANTOM enhancer (32). The proximal convoluted tubule (PCT) had the greatest number of DAR (n=422) followed by the proximal straight tubule (PST), PT_VCAM1, and thick ascending limb (Figure 1C). Less abundant cell types like podocytes and endothelial cells had few if any DAR, which likely reflects our limited power to detect them. Among 1,303 total DAR, 968 were unique because a subset of DAR were shared between multiple cell types (Figure 1C). DAR present in multiple cell types included regions within or near *ATP1B1* and *KCNE1B. ATP1B1* encodes a subunit of the sodium potassium ATPase and *KCNE1B* encodes a subunit of the voltage-gated potassium channel, suggesting that DAR in diabetes may elicit a conserved effect on ion transport across nephron segments.

We grouped DAR from the proximal convoluted tubule (PCT) and proximal straight tubule (PST) and annotated them with the nearest protein coding gene to perform gene ontology enrichment. Genes near proximal tubule DAR were enriched for pathways involved in response to insulin, cellular response to peptides, response to hormone stimuli, and ion transport (Figure 1D). Insulin resistance is a key feature of DKD and we identified proximal tubule DAR with decreased chromatin accessibility near multiple genes that regulate insulin signaling (Supplemental Table 3) (33). There was a DAR in the second intron of the insulin receptor (*INSR*) that showed decreased accessibility in the proximal tubule (Figure 1E, Orange Arrow). This region was predicted to regulate *INSR* expression via a CCAN (Figure 1E, Green Arcs) and was associated with decreased *INSR* expression in the corresponding snRNA-seq dataset (Figure 1F). The loop of Henle is a key regulator of sodium reabsorption where we identified two DAR in the promoter region of *ATP1B1* (Supplemental Table 3). These DAR were present in both the ascending thin limb (ATL) and thick ascending limb (TAL1, TAL2) where they showed decreased chromatin accessibility. Decreased chromatin accessibility was associated with decreased *ATP1B1* expression in the same cell types in the corresponding snRNA-seq dataset. Together, these findings suggest that DKD is associated with changes in chromatin accessibility that regulate expression of genes important for insulin signaling and sodium reabsorption.

We compared the proximal convoluted tubule (PCT) and PT_VCAM1 to identify changes in chromatin accessibility associated with emergence of the pro-inflammatory PT_VCAM1 cell state (Supplemental Table 4). There were 4,498 DAR and the majority showed decreased accessibility (N=3,055, 68%). Gene ontology analysis of nearby protein-coding genes showed enrichment for pathways involved in kidney development, metabolism, amino acid transport, epithelial cell proliferation, response to glucocorticoids, and regulation of transforming growth factor beta signaling. ATAC peaks with increased chromatin accessibility in PT_VCAM1 were located near pro-inflammatory genes like *IL-6, CD40*, and *TGFB2* in addition to genes involved in proliferation like *EGFR* and *MYC*.

### Single Nucleus RNA Sequencing to Detect Differentially Expressed Genes in Type 2 Diabetes

A total of eleven snRNA-seq libraries were aggregated with cellranger (10X Genomics) and analyzed with Seurat following doublet removal and batch effect correction (4,31). The snRNA-seq dataset included six healthy control samples and five with DKD. A total of 39,176 cells passed quality control filters and all major cell types in the kidney cortex were represented (Figure 2A), including the PT_VCAM1 subpopulation (3). snRNA-seq cell types largely expressed the same lineage-specific markers that showed increased chromatin accessibility in the snATAC-seq dataset (Supplemental Figure 3) and were enriched for cell-specific genes (Supplemental Table 5). There was a trend toward greater proportion of PT_VCAM1 in DKD compared to healthy controls (mean proportion 0.06 vs. 0.02, Student’s t-Test p=0.051). We compared individual cell types between healthy control and DKD samples to identify cell-specific differentially expressed genes (Supplemental Table 6). The cell type with the greatest number of differentially expressed genes was the proximal tubule (Figure 2B, N=607, padj < 0.05, |avg_log2FC| > 0.25). Similar to our findings from the snATAC-seq analysis, a subset of differentially expressed genes were shared between multiple cell types (Figure 2C). These shared genes were enriched for pathways involved in regulation of cell growth, cellular response to hypoxia, angiogenesis, cellular response to insulin stimulus, glucocorticoid signaling, and ion transport. For example, *INSR* showed decreased expression in the proximal tubule (Figure 1F), thick ascending limb, and distal convoluted tubule (Supplemental Table 6).

**Figure 2.**
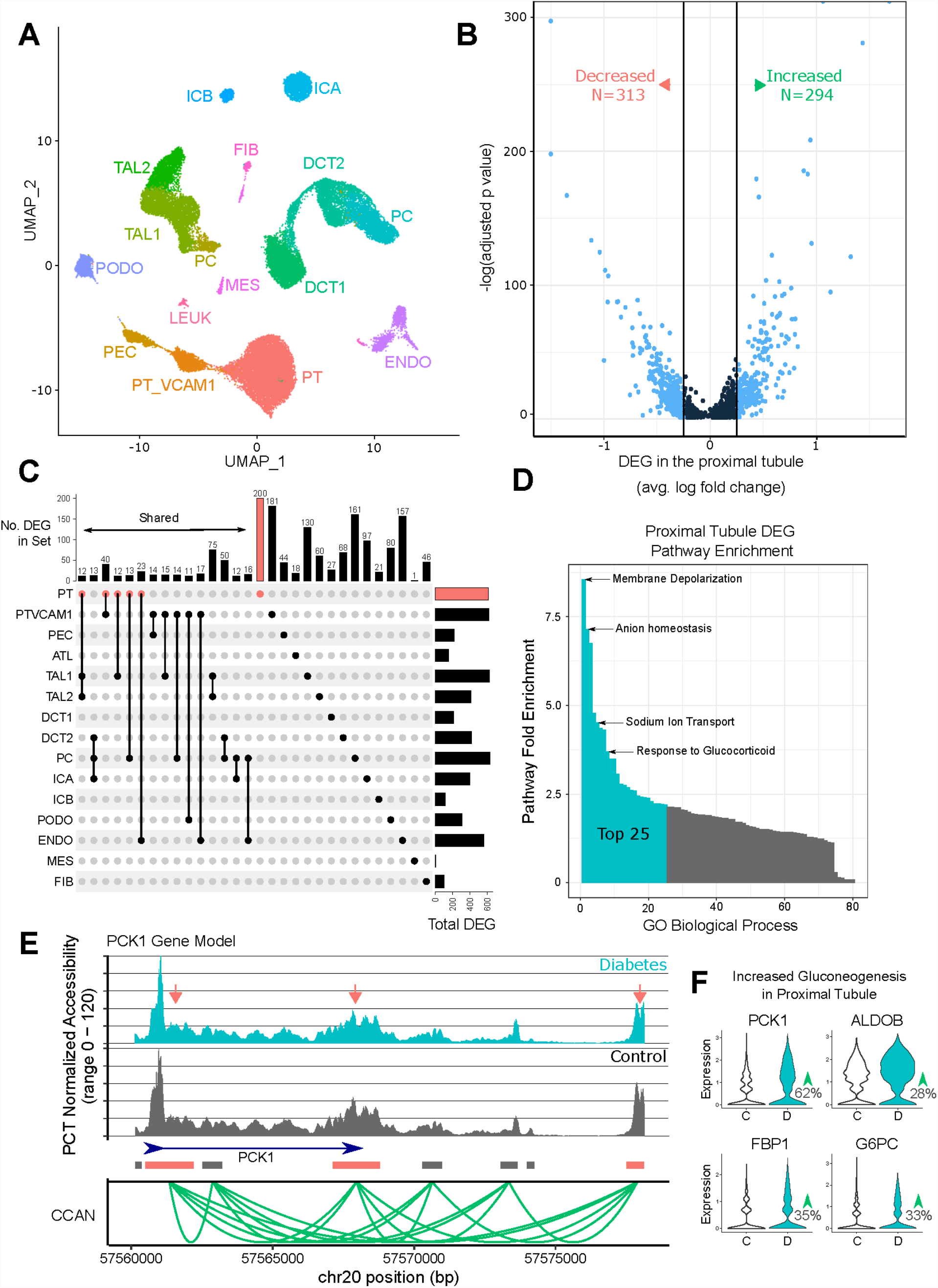
snRNA-seq of human DKD. **A) UMAP embedding of snRNA-seq dataset.** Six healthy control and five DKD samples were aggregated, preprocessed, and filtered. A total of 39,176 cells are depicted. PT-proximal tubule, PT_VCAM1-VCAM1(+) proximal tubule, PEC-parietal epithelial cells, ATL-ascending thin limb, TAL1-CLDN16(-) thick ascending limb, TAL2-CLDN16(+) thick ascending limb, DCT1-early distal convoluted tubule, DCT2-late distal convoluted tubule, PC-principal cells, ICA-type A intercalated cells, ICB-type B intercalated cells, PODO-podocytes, ENDO-endothelial cells, MES-mesangial cells and vascular smooth muscle cells, FIB-fibroblasts, LEUK-leukocytes. **B) Differentially expressed genes in the diabetic proximal tubule**. Healthy control proximal tubule was compared to DKD to identify cell-specific DEG in the snRNA-seq dataset using the Seurat FindMarkers function. DEG were evaluated with a Bonferroni-adjusted p-value for an FDR < 0.05 and significant DEG are displayed. DEG that met an absolute log2-fold-change threshold of 0.25 (vertical bars) are colored light blue. **C) DEG in DKD that are cell-specific or shared between cell types**. DEG that were either shared between multiple cell types or unique to a specific cell type are displayed. DEG shared between multiple cell types are limited to groups that share ten or more DEG. **D) Proximal tubule DEG pathway enrichment**. Significant cell-specific DEG from proximal tubule were used to perform gene ontology enrichment with Panther. Fold-enrichment for all significant GO biological processes is shown and the top 25 are highlighted. **E) Proximal tubule-specific DAR and ATAC peaks in *PCK1***. snATAC-seq coverage plots for DKD and control PCT are displayed in relation to the *PCK1* gene body. The orange arrows indicate multiple DAR that show decreased accessibility in DKD (Supplemental Table 3). Green arcs depict the nodes of a cis-coaccessibility network (CCAN) surrounding the *PCK1* gene body. **F) Proximal tubule shows increased expression of gluconeogenic genes by snRNA-seq**. Healthy control proximal tubule was compared to DKD proximal tubule in the snRNA-seq dataset to identify differentially expressed genes with the FindMarkers function and visualized as violin plots. DKD proximal tubule showed increased expression of *PCK1, ALDOB, FBP1*, and *G6PC* (see Supplemental Table 6 for adjusted p-values).

We examined proximal tubule-specific changes in gene expression to determine if the same pathways we identified by snATAC-seq were present in the snRNA-seq dataset. Gene ontology analysis of differentially expressed genes in the proximal tubule showed significant overlap with snATAC-seq pathways, including membrane depolarization, anion homeostasis, sodium ion transport, and glucocorticoid signaling (Figure 2D). The diabetic proximal tubule showed a modest increase in expression of GR (*NR3C1*, fold-change = 1.14, padj = 4.7 × 10^-10), although it did not meet the log-fold change threshold. The proximal tubule also showed increased expression of sodium glucose cotransporter 2 (SGLT2, fold-change = 1.24, padj = 1.9×10^-30) and increased expression of the rate-limiting enzyme in gluconeogenesis (*PCK1*, fold-change = 1.63, padj = 1.8×10^-41). Comparison with the corresponding snATAC-seq dataset showed multiple proximal tubule DAR near *PCK1* (Figure 2E, Orange Boxes), suggesting that changes in chromatin accessibility may lead to increased *PCK1* expression (Figure 2F). There were no DAR near *SLC5A2* that met the adjusted p-value threshold. The *PCK1* DAR were located both within and distal to its gene body where they interacted with the promoter via a CCAN (Figure 2E, Green Arcs). Additional enzymes in the gluconeogenic pathway were also upregulated in the diabetic proximal tubule (Figure 2F). Together, these findings suggest that the diabetic proximal tubule increases expression of genes that promote both glucose reabsorption (*SLC5A2*) and glucose production (*PCK1, ALDOB, FBP1, G6PC*).

The diabetic thick ascending limb (TAL1) had 622 differentially expressed genes compared to healthy controls (padj < 0.05, |avg_log2FC| > 0.25). The differentially expressed genes in TAL1 were enriched for pathways involved in nitric oxide signaling, ATP biosynthesis, anion transport, and cellular response to cAMP, EGFR signaling, glucocorticoids, hypoxia, and insulin. Similar to the diabetic proximal tubule, there was decreased expression of *INSR* (fold-change = 0.76, padj = 1.5×10^-17) and increased expression of GR (*NR3C1*, fold-change = 1.26, padj = 4.7×10^-10). There was also decreased expression of *HSD11B2* (fold-change = 0.71, padj = 5.3×10^-33), which is the enzyme that catalyzes the conversion of cortisol to the inactive metabolite cortisone to protect nonselective activation of MR. In fact, decreased *HSD11B2* expression was observed in every cell type in the distal nephron (Supplemental Table 6). These data support our hypothesis that the diabetic nephron has increased GR signaling due to increased GR expression and decreased activity of the enzyme responsible for metabolizing cortisol.

We compared the proximal tubule and PT_VCAM1 to identify differentially expressed genes associated with the PT_VCAM1 cell state. There were 3,842 differentially expressed genes (Supplemental Table 7) enriched for pathways involved in cell migration, EGFR signaling, insulin receptor signaling, histone deacetylation, regulation of glycolysis, and TGF-beta signaling. For example, *INSR* expression was decreased in PT_VCAM1 relative to PT (fold-change = 0.67, padj = 4.1×10^-53) and *TGFBR2* was increased (fold-change = 1.34, padj = 5.4×10^-34). These changes were accompanied by a modest increase in GR expression (*NR3C1*, fold-change = 1.07, padj = 2.0×10^-10) and a marked reduction in *FKBP5* (fold-change = 0.45, padj = 4.9×10^-117).

### Cell-specific Transcribed Cis-Regulatory Elements

Transcribed cis-regulatory elements (tCRE) confer cell type specificity and the majority are located in enhancer and promoter regions where they overlap with ATAC peaks (34,35). Transcriptional start sites (TSS) can be identified by 5’ RNA sequencing if read 1 is long enough (> 81bp) to include the junction between the template switch oligo (TSO) and TSS. This type of analysis is compatible with single cell 5’ paired-end chemistry (SC5P-PE, 10X Genomics), but will not work with libraries that only use read 2 for alignment (SC5P-R2, 10X Genomics). We analyzed two healthy control and two DKD snRNA-seq libraries with SC5P-PE sequencing to identify *de novo* transcriptional start sites in CRE using Single Cell Analysis of Five-prime Ends (SCAFE) (36,37). SCAFE analyzes the 5’ end of RNA transcripts to identify reads mapping to the junction between the TSO and cDNA sequence to localize TSS within tCRE after filtering false positives with a logistic regression classifier. We identified 37,698 tCRE across all cell types (Figure 3A, Supplemental Table 8). The majority of tCRE were near a protein-coding TSS (mean distance = 3823 +/-43,885), but there was a significant proportion of tCRE in intronic (Figure 3B, 11847/37,698, 31%) and intergenic regions (Figure 3B, 1367/37,698, 3%). Some of these tCRE may represent enhancer RNA (eRNA), which are a family of non-coding RNA that regulate gene expression and enhancer activity in a cell-specific manner (38). The majority of tCRE were overlapping with a snATAC-seq peak (Figure 3C, 23,048/37,698, 61%) and approximately half of snATAC-seq DAR were overlapping with a tCRE (494/968, 51%, hypergeometric test p=3.6×10^-5). A small minority of tCRE were cell-type-specific (N=361/37,698, 1%, Supplemental Table 7), but corresponded to well-known cell-type-specific genes. For example, there was a cell-specific tCRE in podocytes in the promoter region of *NPHS1* and a cell-specific tCRE in the proximal tubule in the promoter region of *CUBN* (Supplemental Table 7). These data suggest that 5’ snRNA-seq datasets contain complementary information that can be used to evaluate the activity of CRE identified by snATAC-seq.

**Figure 3.**
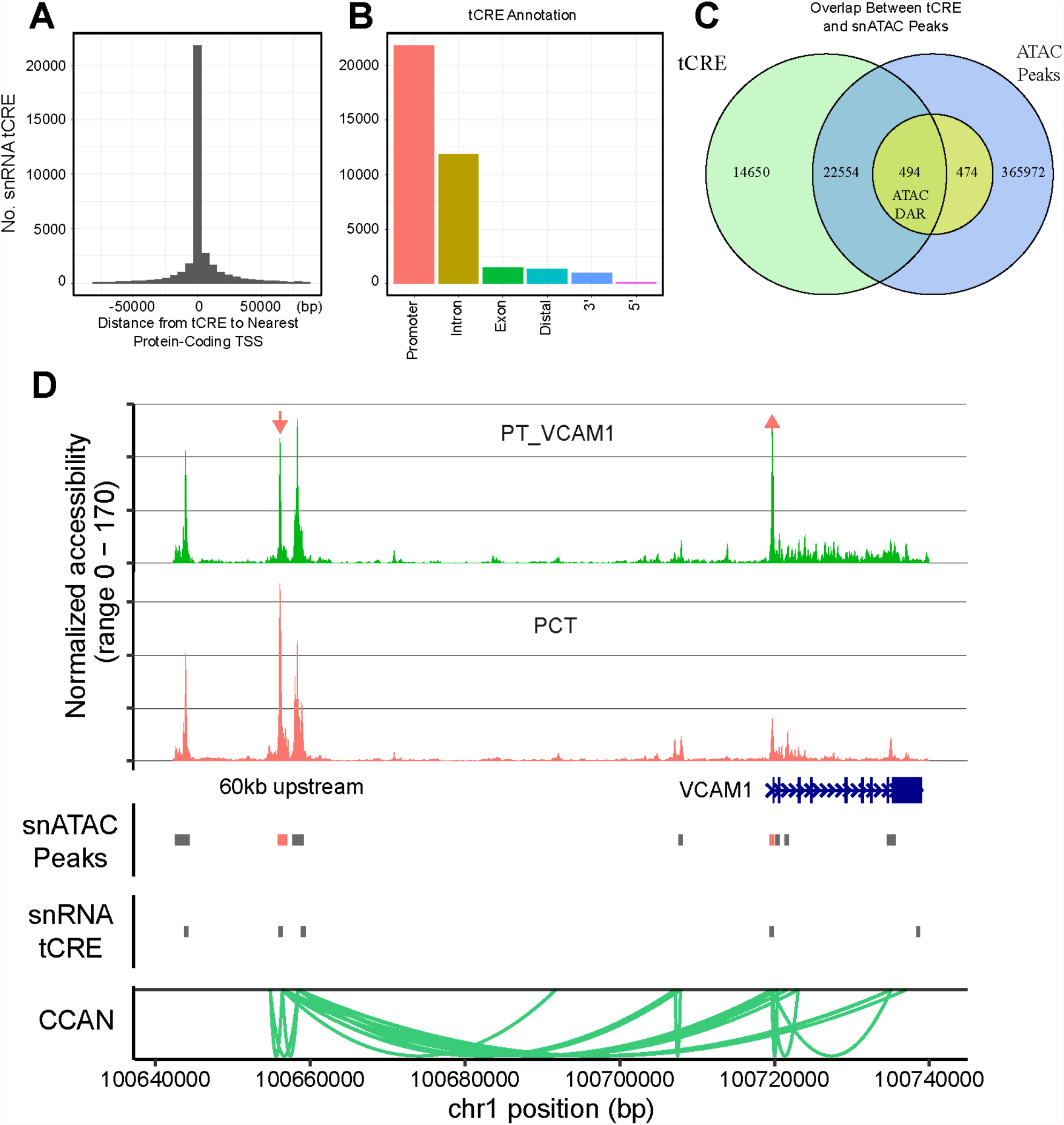
Transcribed cis-regulatory elements (tCRE) detected by 5-prime snRNA-seq. **A) Distance from tCRE to TSS.** Two healthy control and two DKD samples were sequenced with 5’ paired-end sequencing and analyzed with SCAFE. A total of 37,698 tCRE were annotated with ChIPSeeker and displayed relative to the nearest TSS. **B) Annotation of tCRE**. The relative proportion of tCRE in promoters, introns, exons, distal intergenic, 3-prime, and 5-prime regions is shown. **C) Overlap between tCRE and ATAC peaks**. tCRE were intersected with cell-specific ATAC peaks and DAR in DKD using GenomicRanges. **D) Proximal convoluted tubule and PT_VCAM1 DAR and ATAC peaks in *VCAM1***. snATAC-seq coverage plots for PT_VCAM1 (green) and PCT (orange) are displayed in relation to the *VCAM1* gene body. The orange arrows indicate DAR that show either increased or decreased accessibility in PT_VCAM1 relative to PCT (Supplemental Table 4). snATAC-seq peaks accessible in the proximal tubule are displayed (snATAC peaks, gray boxes) in the same track as PCT DAR (snATAC peaks, orange boxes). snRNA-seq tCRE regions are displayed below snATAC-seq peaks and DAR (snRNA tCRE, gray boxes). Green arcs depict the nodes of a cis-coaccessibility network (CCAN) surrounding the *VCAM1* gene body.

We compared healthy control to DKD samples to identify cell-specific differential tCRE in diabetes. Across all cell types, we detected a total of 293 differential tCRE (Supplemental Table 9). These tCRE included 139 unique regions, which were enriched for pathways involved in mitochondrial electron transport and angiogenesis. Multiple cell types showed increased transcription of CRE in promoters associated with oxidative phosphorylation like *MT-CO1, MT-CO2* and *MT-CO3*. We also compared the proximal tubule to PT_VCAM1 to identify tCRE that are enriched in the PT_VCAM1 cell state (Supplemental Table 10). Among 204 differential tCRE in PT_VCAM1, one of the most enriched tCRE was the *VCAM1* promoter (fold change = 1.46, padj = 1.1×10^-82). The *VCAM1* promoter tCRE showed increased chromatin accessibility in the corresponding snATAC dataset (Figure 3D) and was associated with two additional tCRE located approximately 60kb upstream. Each of the upstream tCRE were near a snATAC-seq peak and linked to the *VCAM1* promoter via a CCAN (Figure 3D). We previously reported that this upstream CRE binds NFkB by chromatin immunoprecipitation PCR (3). NFkB signaling induces *VCAM1* expression in the proximal tubule, which raises the possibility that NFkB binding to the upstream CRE is also associated with transcription of enhancer RNA (38,39). Together, these data suggest that single cell analysis of 5’ ends may help to identify enhancers by prioritizing CRE that are actively transcribed.

### Glucocorticoid Receptor CUT&RUN in Bulk Kidney Cortex

Cellular response to glucocorticoids is influenced by pre-existing chromatin accessibility state where the majority of GR binding sites localize to open chromatin regions (40,41). We used cleavage under targets and release using nuclease (CUT&RUN) to directly measure GR binding in bulk kidney cortex obtained from a healthy donor (42). We identified 4,362 GR binding sites (Supplemental Table 11) located in promoter regions (N=2889, 66%), introns (N=744, 17%) and distal intergenic regions (N=567, 13%). The density of cell-specific ATAC peaks closely-resembled the density of GR CUT&RUN sites across the genome, which suggests that GR predominantly binds to areas of open chromatin in the kidney (Figure 4A). GR binding sites overlapped with cell-specific ATAC peaks (N=3066, 70%); many of which were shared between multiple cell types. We visualized the intersection between cell-specific ATAC peaks and CUT&RUN sites to identify individual cell types or groups of cells that share ten or more GR binding sites (Figure 4B). The presence of GR binding sites within cell-specific ATAC peaks suggests GR signaling is controlled by chromatin accessibility and regulated by distinct GR modules shared across cell types. For example, there were GR binding sites in ATAC peaks unique to the proximal tubule (N=64, Figure 4B), unique to the distal nephron (N=15, Figure 4B), and shared between the proximal tubule and distal nephron (N=60, Figure 4B). Similarly, there were GR binding sites unique to lymphocytes and shared between the proximal tubule and lymphocytes.

**Figure 4.**
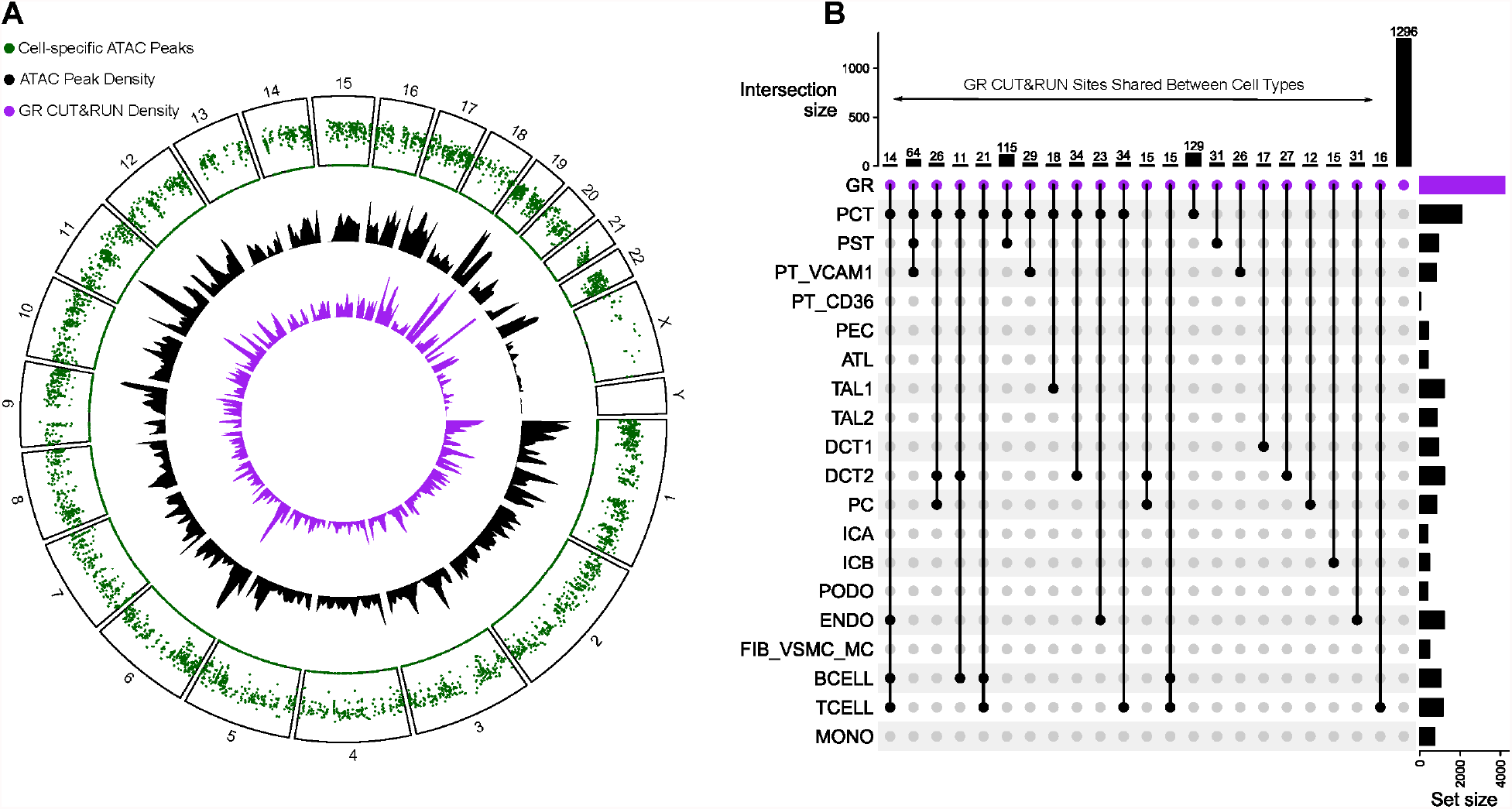
Glucocorticoid receptor (GR) CUT&RUN in bulk kidney cortex. **A) Density of GR CUT&RUN sites relative to cell-specific ATAC peaks.** Cell-specific ATAC peaks were identified with the Seurat FindMarkers function (Supplemental Table 2) and converted into a rainfall plot (green track) using the circlize package in R. Each dot in the rainfall plot corresponds to a cell-specific ATAC peak and the y-axis corresponds to the minimal distance between the peak and its two neighboring regions. Clusters of regions appear as a “rainfall” in the plot. The density of cell-specific ATAC peaks (black track) and GR CUT&RUN peaks (purple track) are shown adjacent to the rainfall plot using a 10Mb default window size. **B) Upset plot showing intersection between GR CUT&RUN sites and cell-specific ATAC peaks**. Cell-specific ATAC peaks that contain a bulk kidney GR CUT&RUN site that are also shared between multiple cell types are displayed. Only intersections with ten or more GR CUT&RUN sites are included in the plot. GR CUT&RUN sites that do not intersect a cell-specific ATAC peak are displayed as the black bar to the far right (N=1,296).

### Transcription factor motif enrichment and activity in the diabetic nephron

We used the JASPAR database to identify over-represented transcription factor motifs in cell-specific ATAC peaks and DAR in DKD (43). Cell-specific ATAC peaks were enriched for established transcription factors that drive cell type differentiation like HNF4A in the proximal tubule and TFAP2B in the distal nephron (Supplemental Table 12). Transcription factors that were enriched in DAR provide insight into cell-specific signaling pathways that are altered in DKD. PCT DAR were significantly enriched for NR3C1 and NR3C2 motifs (Supplemental Table 13, NR3C1 fold enrichment = 2.1, p = 3.8×10^-259; NR3C2 fold enrichment=2.1, p = 2.4×10^-288). NR3C1 is the canonical binding motif for GR and NR3C2 is the binding motif for MR. The presence of NR3C1 and NR3C2 motifs within PCT DAR suggests that chromatin accessibility may regulate corticosteroid signaling in the diabetic proximal tubule (44,45). We also saw enrichment of KLF9 and FOXO3 motifs within PCT DAR, which are downstream of GR activation (Supplemental Table 13). One of the most enriched motifs in PCT DAR was HINFP (fold enrichment = 6.9, p = 6.3×10^-308). Histone H4 transcription factor (HINFP) interacts with a component of the MeCP1 histone deacetylase complex (HDAC) involved in transcriptional repression, which may explain why the majority of DAR showed decreased chromatin accessibility (46). To help prioritize active signaling pathways in DKD, we identified transcription factor motifs that were both differentially expressed and enriched in DAR (Figure 5A). HIF1A showed increased expression and was enriched in DAR in multiple distal nephron cell types (PC, ICA, DCT2) and PT_VCAM1. HIF1A is hypoxia inducible factor 1 subunit alpha, a master regulator of cellular response to hypoxia in the kidney (47). GR showed increased expression in the proximal tubule (PT) and thick ascending limb (TAL1) where NR3C1 motifs were also enriched in DAR. In contrast, MR showed decreased expression in the distal nephron (PC, DCT2), but increased expression in the thick ascending limb (TAL1). Together, these data suggest that corticosteroid signaling is altered in the diabetic proximal tubule and thick ascending limb where multiple cell types may be exposed to a hypoxic environment.

**Figure 5.**
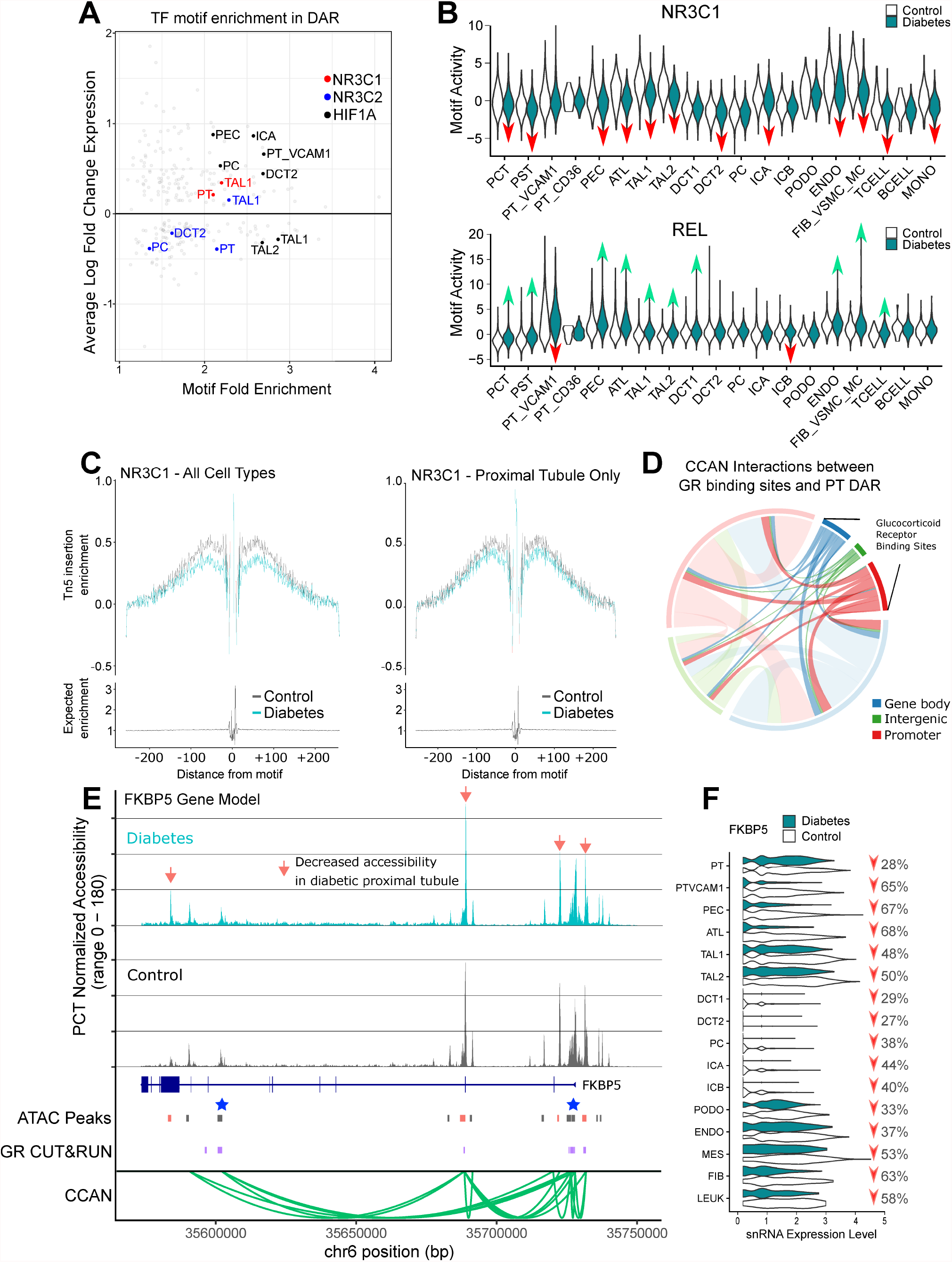
Glucocorticoid receptor (GR) binding and *FKBP5* in DKD. **A) Cell-specific transcription factor expression and motif enrichment in DKD.** Transcription factors that were both differentially expressed (Supplemental Table 6) and showed motif enrichment in cell-specific DAR (Supplemental Table 13) were visualized in a scatter plot. Cell types that showed differential expression and motif enrichment for GR (NR3C1, red), MR (NR3C2, blue), and HIF1A (HIF1A, black) motifs are highlighted with different colors. PT-proximal tubule, PT_VCAM1-*VCAM1*(*+*) proximal tubule cells, PEC-parietal epithelial cells, TAL1-*CLDN16*(-) thick ascending limb, TAL2-*CLDN16*(+) thick ascending limb, DCT2-late distal convoluted tubule, PC-principal cells, ICA-type A intercalated cells. **B) Cell-specific chromVAR motif activity for GR and REL**. chromVAR was used to compute cell-specific activities for NR3C1 (GR) and REL motifs for control and DKD cell types. Red arrows indicate significantly decreased motif activity and green arrows indicate significantly increased motif activity (see Supplemental Table 14 for average log-fold-change and adjusted p-values). **C) Transcription factor footprinting for GR**. Transcription factor footprinting analysis was performed for NR3C1 (GR) for all cell types and for PCT only to quantitate Tn5 insertion enrichment in healthy control and DKD using Signac. **D) Interaction between PCT DAR and hTERT-RPTEC GR CUT&RUN sites**. PCT DAR in DKD (Supplemental Table 3) were intersected with cis-coaccessibility networks (CCAN) to identify all CCAN links that contain at least one PCT DAR. These regions were intersected with hTERT-RPTEC GR CUT&RUN sites and visualized with the circlize package in R to identify links between GR CUT&RUN sites and PCT DAR. Sites were annotated with ChIPSeeker. **E) PCT-specific DAR and ATAC peaks in *FKBP5***. snATAC-seq coverage plots for DKD and control PCT are displayed in relation to the *FKBP5* gene body. The orange arrows indicate multiple DAR that show decreased accessibility in DKD (Supplemental Table 3). PCT-specific ATAC peaks (Peaks, dark gray boxes) and DAR (Peaks, orange boxes) are shown in relation to hTERT-RPTEC CUT&RUN sites (GR, purple boxes) and a cis-coaccessibility network (CCAN, green arcs) surrounding the *FKBP5* gene body. Blue stars indicate sites targeted by CRISPRi. **F) Cell-specific expression of *FKBP5* by snRNA-seq**. Individual cell types were compared between control and DKD with the Seurat FindMarkers function and visualized to display relative change in *FKBP5* expression. Red arrows indicate significantly decreased *FKBP5* expression (see Supplemental Table 6 for adjusted p-values).

We used chromVAR to compare transcription factor activity between healthy control and diabetic cell types (Supplemental Table 14). chromVAR is a tool for inferring transcription-factor-associated chromatin accessibility in single cells that helps to address sparsity inherent in snATAC-seq datasets (48). This is a qualitatively different and unbiased approach when compared to transcription factor motif enrichment analysis because it is not limited to a pre-specified list of cell-specific DAR. Diabetic PCT showed decreased transcription factor activity for NR3C1 (Figure 5B, fold-change = 0.56, padj = 1.1×10^-70) and increased activity for REL motifs (Figure 5B, fold-change = 2.08, padj = 6.9×10^-212), which was a pattern observed throughout the nephron. These data support our transcription factor motif enrichment analysis by further demonstrating that NR3C1 motifs localize to areas of decreased chromatin accessibility in diabetes and REL motifs localize to areas of increased accessibility.

### Glucocorticoid Receptor Footprinting with snATAC-seq and CUT&RUN in RPTEC

We used the snATAC-seq dataset to perform transcription factor footprinting for GR to visualize the relationship between NR3C1 motifs and chromatin accessibility. Across all cell types, there was a well-defined footprint immediately surrounding NR3C1 motifs (Figure 5C). DKD samples showed reduced chromatin accessibility surrounding NR3C1 motifs, however, this effect was attenuated when we limited our analysis to the proximal tubule (Figure 5C). We cultured immortalized renal proximal tubule epithelial cells (hTERT-RPTEC, ATCC) and performed CUT&RUN to find 22,539 consensus GR binding sites that were not present in IgG-stimulated negative control samples (Supplemental Table 15). hTERT-RPTEC media is supplemented with 25ng/ml hydrocortisone (ie. cortisol) and is a model of long-term glucocorticoid exposure. Nearly half of PCT DAR (168/422, 39%) were overlapping with a GR CUT&RUN site. These findings are comparable to the chromVAR analysis, which showed that approximately 47% of healthy control PCT and 35% of diabetic PCT ATAC peaks contain an NR3C1 motif. GR binding sites that do not directly overlap a PCT DAR may interact with DAR via CCAN (Figure 5D). For example, we identified multiple GR binding sites throughout the *FKBP5* gene body located in promoter and intronic regions (Figure 5E, Purple Boxes). Some of the GR binding sites in *FKBP5* co-localized with PCT DAR (Figure 5E, Orange Boxes), but others did not. *FKBP5* expression was decreased throughout the entire nephron (Figure 5F), which highlights its potential importance in DKD and raises the possibility that changes in chromatin accessibility regulate its expression. The high proportion of overlap between GR binding sites and snATAC-seq PCT DAR was especially striking given that hTERT-RPTEC are a cell culture model that does not fully recapitulate the normal proximal tubule. We profiled open chromatin regions in hTERT-RPTEC and primary RPTEC using Omni-ATAC and compared them to the PCT snATAC-seq dataset. Approximately 59% of hTERT-RPTEC ATAC peaks (N=57,675/96,162, Supplemental Table 16) and 56% of primary RPTEC ATAC peaks (N=80,322/141,198, Supplemental Table 17) were overlapping with cell-specific PCT snATAC-seq peaks. These data suggest that hTERT-RPTEC and primary RPTEC capture roughly half of the chromatin accessibility profile of a normal proximal tubule cell.

### Validation of Differentially Expressed Genes in a Bulk RNA-seq Dataset of Human DKD

We analyzed a previously-published bulk RNA-seq dataset of human DKD to determine if our snRNA-seq findings are broadly generalizable. The dataset published by Fan et. al consisted of 9 controls, 6 with early DKD, and 22 with advanced DKD (49). Early DKD was defined as eGFR > 90mL/min/1.73m^2 and UACR < 300mg/g. Advanced DKD was defined as eGFR < 90mL/min/1.73m^2 or UACR > 300mg/g. According to this definition, samples from our study would be categorized as advanced DKD because all of them had either eGFR < 90mL/min/1.73m^2 or proteinuria. Another important difference between our study and the study by Fan et al. is that mean eGFR of control samples from our study (66 +/-25 ml/min/1.73m^2) was significantly less than eGFR of control samples from their study (87 +/-9.8 ml/min/1.73m^2, p=0.03).

We compared the transcriptional profile of advanced DKD to control samples from Fan et al. to identify 9,632 differentially expressed genes (Supplemental Table 18A, BH padj < 0.05). Roughly half of these differentially expressed genes were upregulated (N=5,181) and the remaining were downregulated (N=4,451). These differentially expressed genes were enriched for familiar pathways including amino acid metabolism, B cell receptor signaling, T cell differentiation, response to tumor necrosis factor, response to peptide hormone, cellular response to hormone stimulus, and ion transmembrane transport. The enrichment of pathways involved in lymphocyte signaling and differentiation likely reflects the greater proportion of leukocytes present in DKD samples compared to control samples. We previously demonstrated that advanced DKD samples from Fan et al. contain an increased proportion of leukocytes and PT_VCAM1 (3). Advanced DKD samples showed increased expression of GR (*NR3C1*, fold-change = 1.16, padj = 0.02) and *VCAM1* (fold-change = 1.36, padj = 0.006) and decreased expression of *INSR* (fold-change = 0.49, padj = 1.6×10^-12), *HSD11B2* (fold-change = 0.38, padj = 3.6×10^-8) and *FKBP5* (fold-change = 0.46, padj = 0.0009). Next, we compared early DKD samples to controls to identify 1,041 differentially expressed genes, among which 385 were upregulated and 656 were downregulated (Supplemental Table 18B). The differentially expressed genes in early DKD were enriched for pathways involved in response to epidermal growth factor, cellular response to glucocorticoids, cellular response to insulin stimulus, and cellular response to tumor necrosis factor. In contrast to the advanced DKD samples, we did not detect differential expression of *NR3C1, INSR, HSD11B2*, or *FKBP5* in early DKD samples. This difference may reflect reduced specificity of bulk RNA-seq and a limited number of early DKD samples (n=6) vs. advanced DKD samples (n=22), or alternatively, that these genes are associated with DKD progression.

### CRISPRi knockdown of FKBP5 Cis-regulatory Elements

We selected two GR binding sites in the *FKBP5* gene body (Figure 5E, Blue Stars) that were located at nodes within a CCAN (Figure 5E, Green Arcs) to target with CRISPRi. Catalytically inactive dCas9 fused to the Krüppel-associated box (KRAB) repression domain (dCas9-KRAB) reduces chromatin accessibility to induce targeted gene silencing. We transduced primary RPTEC with sgRNA targeting the TSS or intronic CRE in *FKBP5* to repress chromatin accessibility with dCas9-KRAB (Figure 6A) (50). Transduction of sgRNAs targeting the TSS or intronic region induced a 30-50% reduction in *FKBP5* expression compared to non-targeting control sgRNA (Figure 6B). Furthermore, gene silencing was specific to *FKBP5* because CRISPRi did not affect expression of neighboring genes expressed in primary RPTEC (Figure 6C).

**Figure 6.**
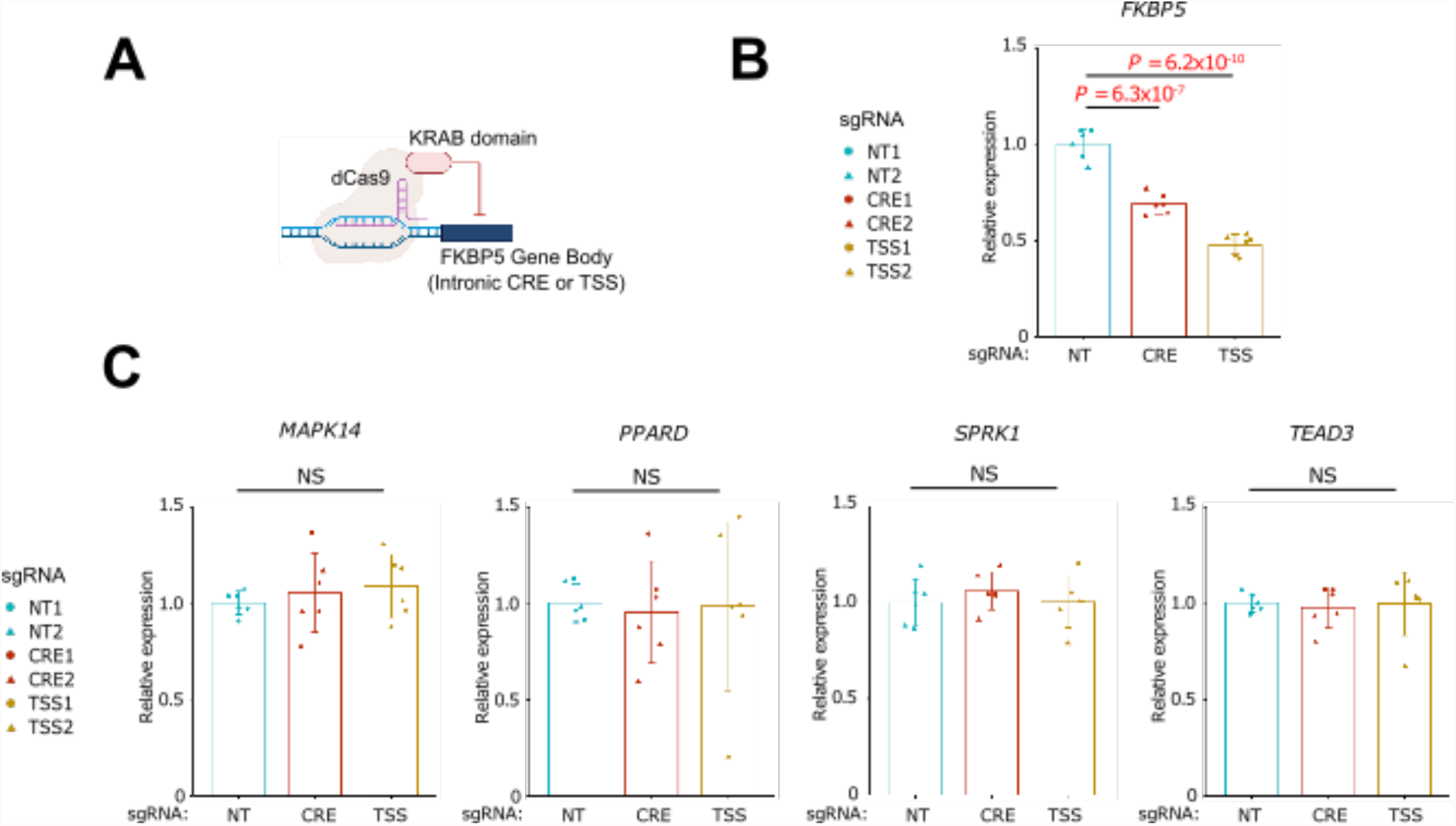
Knockdown of *FKBP5* cis-regulatory elements with CRISPR interference. **A) CRISPR interference diagram.** dCas9-KRAB domain fusion protein and small guide RNAs (sgRNA) were used to target the TSS and a potential intronic CRE in the *FKBP5* gene. Targeted regions are depicted as blue stars in the *FKBP5* gene model diagram in Figure 5. sgRNA primers and region coordinates are provided in Supplemental Table 20. **B) Quantitative PCR of *FKBP5* CRISPRi**. RT and real-time PCR analysis of mRNAs for *FKBP5* and surrounding genes (*MAPK14, PPARD, SPRK1* and *TEAD3*) in primary renal proximal tubular epithelial cells (primary RPTEC) with CRISPR interference targeting the TSS and predicted cis-regulatory element (CRE) for *FKBP5*. NT, non-targeting control. Each group consists of n = 6 data (2 sgRNAs with 3 biological replicates). Bar graphs represent the mean and error bars are the s.d. p-values are calculated with one-way ANOVA and a post-hoc Dunnett’s test for multiple comparisons.

### Partitioned Heritability of Cell-specific ATAC Peaks and Differentially Accessible Regions for Kidney-Function-Related GWAS Traits

GWAS have shown that a growing list of kidney-related traits have a genetic component (51– 54). We downloaded GWAS summary statistics for eGFR, CKD, microalbuminuria, and urinary sodium excretion to determine whether cell-specific chromatin accessibility patterns explain heritability of these traits. First, we partitioned heritability of cell-specific ATAC peaks with stratified linkage disequilibrium score regression to prioritize which cell types explain heritability of kidney-function-related traits after controlling for baseline enrichment (55). The cell types that showed greatest enrichment for heritability of eGFR after correction for multiple comparisons were segments of the proximal tubule (PCT, PST) and the PT_VCAM1 subpopulation (Figure 7A). This relationship between proximal tubule and heritability of eGFR has been previously-described (56). Interestingly, PT_VCAM1 cell-specific peaks also showed increased heritability for CKD, which raises the possibility that genetic background may influence the transition from healthy proximal tubule to PT_VCAM1 (Figure 7A). Multiple segments of the thick ascending limb (TAL1, TAL2) and principal cells (PC) showed enrichment for urinary sodium excretion, which is consistent with their known roles in sodium reabsorption. In contrast, we did not identify any cell types that showed increased heritability for microalbuminuria. This may reflect our reduced sensitivity to detect podocyte-specific ATAC peaks that may regulate this phenotype. Next, we partitioned heritability of cell-specific DAR that change in DKD. Similar to the findings from our cell-specific ATAC peak analysis, the DAR in the proximal tubule showed increased heritability of eGFR (Figure 7B). Furthermore, DAR in the thick ascending limb (TAL1) showed increased heritability of urine sodium excretion (Figure 7B). These data suggest that DKD induces changes in chromatin accessibility in some of the same regions that predict heritability of cell-specific kidney functions. It also raises the possibility that genetic background may modulate chromatin accessibility patterns to influence changes in eGFR or sodium excretion in DKD.

**Figure 7.**
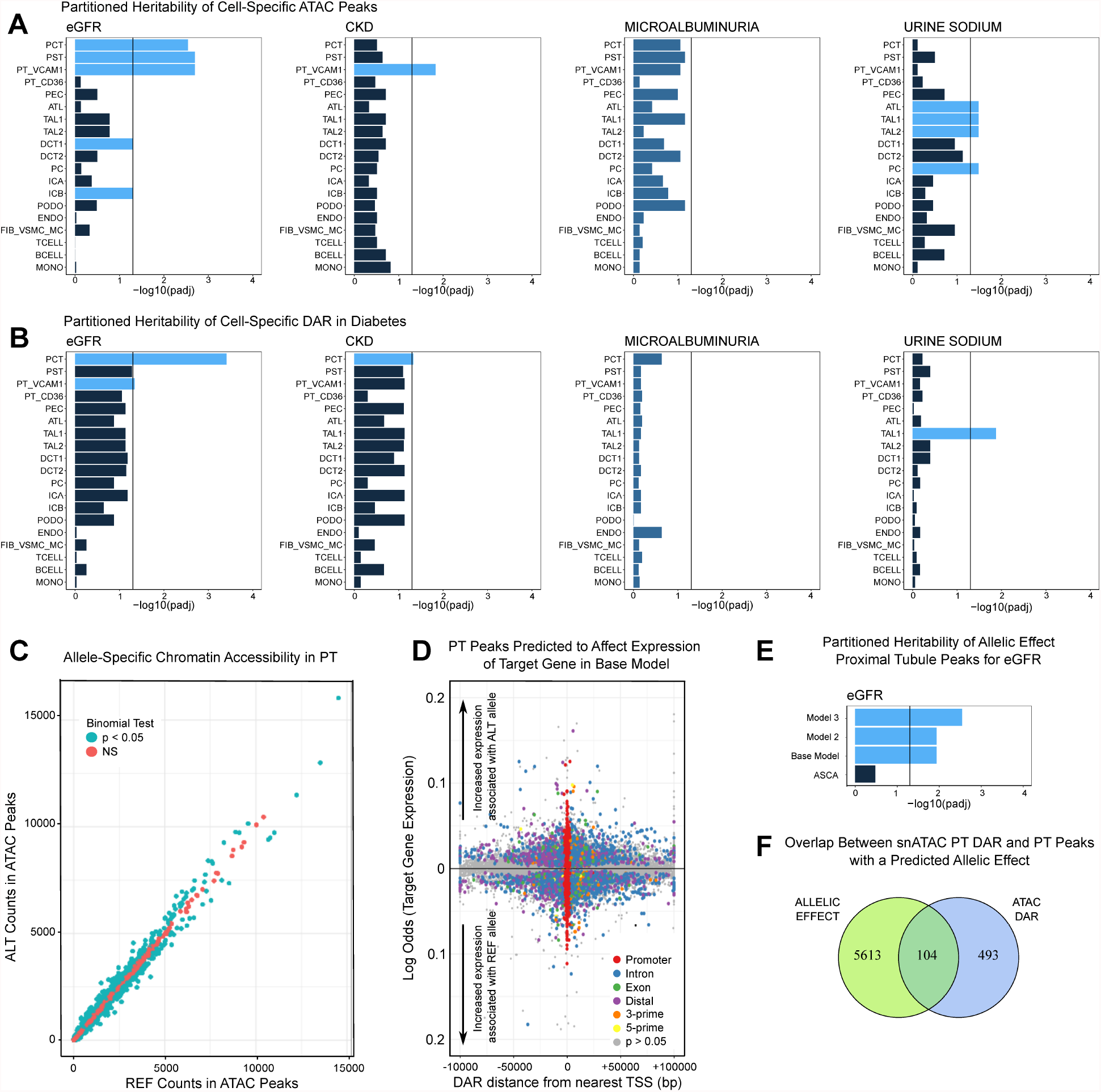
Cell-specific partitioned heritability of GWAS traits and prediction of allelic effects with SALSA. **A) Partitioned heritability of cell-specific ATAC peaks for eGFR, CKD, microalbuminuria, and urinary sodium excretion.** Cell-specific ATAC peaks were identified with the Seurat FindMarkers function (Supplemental Table 2) and prepared as bed files to create custom annotations and generate linkage disequilibrium scores with ldsc. Partitioned heritability was performed using publicly-available GWAS summary statistics and the cell-type-specific workflow. Significance was assessed with an adjusted p-value threshold < 0.05 (vertical bars). **B) Partitioned heritability of cell-specific DAR for eGFR, CKD, microalbuminuria, and urinary sodium excretion in DKD**. Cell-specific DAR were identified by comparing healthy control and DKD cell types with the Seurat FindMarkers function (Supplemental Table 3). Significant regions were prepared as bed files to create custom annotations and generate linkage disequilibrium scores with ldsc. Partitioned heritability was performed using publicly-available GWAS summary statistics and the cell-type-specific workflow. Significance was assessed with an adjusted p-value threshold < 0.05 (vertical bars). **C) Ratio of snATAC-seq fragments mapping to reference and alternate alleles in the proximal tubule**. SALSA was used to identify phased heterozygous SNV and quantitate single cell snATAC-seq fragments in the proximal tubule (PCT, PST) mapping to the reference or alternate allele. Counts were aggregated across all libraries (N=6 control, N=5 DKD) and evaluated for allele specific chromatin accessibility using a binomial test. **D) Predicting an allele-specific effect with SALSA in the proximal tubule**. Proximal tubule snATAC-seq peaks containing a heterozygous SNV were analyzed with the LinkPeaks function in Signac to identify one or more gene targets. Gene target expression was estimated using label transfer from the snRNA-seq to the snATAC-seq object. The presence of a fragment mapping to an alternate allele in a proximal tubule ATAC peak (binary dependent variable) was modeled as a function of target gene expression (continuous predictor variable) after controlling for sample-to-sample variability by including a mixed effect per library with the glmer package. The effect size for each peak is displayed in log-odds where a 1 unit increase corresponds to a 1% increase in target gene expression. Peaks are displayed relative to the nearest TSS and peaks that met the significance threshold (Wald p < 0.05) were annotated with ChIPSeeker to indicate a predicted region (Promoter=red, Intron=blue, Exon=green, Distal intergenic=purple, 3-prime=orange, 5-prime=yellow). Peaks that did not meet the significance threshold are plotted in the bottom layer and colored gray (Wald p > 0.05). **E) Partitioned heritability of proximal tubule peaks with a predicted allelic effect**. Peaks that were associated with changes in target gene expression were partitioned for heritability of eGFR for each of three GLMM evaluated in addition to peaks that met the binomial threshold for allele-specific chromatin accessibility (ASCA, panel C). Significance was assessed with an adjusted p-value threshold < 0.05 (vertical bar). **F) Overlap between proximal tubule DAR and peaks with a predicted allelic effect**. DAR from PCT and PST were intersected with proximal tubule peaks with a predicted effect in Model 3.

### Allele-specific chromatin accessibility as a modifier of gene expression

We created an open-source and containerized workflow for single cell allele-specific analysis called SALSA (https://github.com/p4rkerw/SALSA). SALSA is a tool for genotyping, phasing, mapping bias correction, and modeling of single cell allele-specific counts obtained from snRNA-seq or snATAC-seq datasets. The SALSA workflow includes user-friendly tutorials and is executed in a publicly-available Docker container built on top of the Genome Analysis Toolkit (GATK) developed by the Broad Institute (57). SALSA uses GATK best practices for germline short-variant discovery to identify SNV and indels, which are phased using shapeit4 and a population-based reference from 1000 Genomes (58,59). Phased variants present in the population-based reference are used to perform mapping bias correction and eliminate technical artifacts with WASP (60). Heterozygous germline SNV that overlap ATAC peaks identify single cell allele-specific peak fragments that map to either the reference or alternate allele. In this manner, heterozygous SNV in ATAC peaks are used as markers to assign a peak fragment to one haplotype or the other. This is an attractive approach because the reference haplotype can serve as a perfectly-matched internal control for each individual. We quantitated single cell allele-specific peak fragments in the proximal tubule and plotted the aggregate ratio of fragments mapping to reference or alternate alleles among 43,479 peaks containing a heterozygous SNV (Figure 7C). The majority of peaks had an equal proportion of fragments mapping to each allele (N=35,019, 80%), but a minority of peaks had allelic bias as evaluated by a binomial test (N=8,460, 20%). A significantly smaller proportion of peaks met the adjusted p-value threshold (N=542, 1.2%), suggesting that most proximal tubule peaks do not show allelic bias when aggregated across a population. Next, we integrated snRNA-seq and snATAC-seq datasets to apply an algorithm developed by Ma et al. to identify ATAC peaks that are correlated with expression of nearby genes after correction for distance, GC content, peak accessibility, and peak width (61). This approach helped to identify one or more gene targets for each peak containing a heterozygous SNV. We developed a simple mixed effect logistic regression model where the binary dependent variable was coded as the presence of an alternate allele in an ATAC peak fragment and the continuous predictor variable was single cell target gene expression in the integrated multimodal dataset. A mixed effect per sample was included to control for pseudo-replication bias (62). Our approach is a modification of a previously-published model in SnapATAC used to identify gene-enhancer pairs that coded the dependent variable as ‘open’ or ‘closed’ and omitted the mixed effect (63). Our base model evaluates whether increased or decreased expression of a target gene is predictive of the presence of an alternate allele within an ATAC peak. In the simplest terms, we can ask if the presence of a SNV in an ATAC peak is associated with changes in gene expression.

In the base model, we evaluated 66,828 peak-gene combinations to estimate the effect of gene expression on the presence of a heterozygous SNV in an ATAC peak (Figure 7D, Supplemental Table 19A). The peak-gene combinations included 42,990 unique ATAC peaks where the majority had either one (N=28,989, 67%) or two gene targets (N=8,767, 20%). Approximately 11% of peak-gene combinations showed evidence of an allele-specific effect (Figure 7D, 7512/66,828, Wald test p < 0.05), which decreased to 1% after adjustment for multiple comparisons (N=714, 1%). There were 5,908 unique ATAC peaks with at least one significant peak-gene allelic effect, predominantly in promoter (N=2,312, 39%) and intronic regions (N=2,082, 35%). The number of peak-gene combinations that showed increased expression in association with an alternate allele (N=3,664, 48%) was similar to the number of combinations that showed increased expression in association with the reference allele (N=3,848, 52%). Among peaks that met the significance threshold, the median absolute coefficient value was 0.01 (log odds) for a 1% increase in target gene expression. For a 10% increase in gene expression, this translates to the typical ATAC peak being 1.10 times more likely to contain an alternate allele in the base model. In subsequent models, we added a fixed effect for diabetes (Model 2) and a fixed effect for diabetes with an interaction term between target gene expression and diabetes (Model 3). Model 2 had 7,557 peak-gene combinations where expression was a significant predictor of the presence of an alternate allele (Supplemental Table 19B, Wald test p < 0.05), which included 6,980 that were also identified in the base model. Since we know that diabetes can alter gene expression, Model 3 included an interaction term between expression and diabetes. Model 3 had 7,353 peak-gene combinations where expression was a significant predictor of the presence of an alternate allele (Supplemental Table 19C, Wald test p < 0.05), which included 4,577 that were also identified in the base model. Approximately 28% of these peak-gene combinations (N=2,097/7,353) had a significant interaction between expression and diabetes. Additionally, diabetes was a significant predictor of the presence of an alternate allele in 20% of peak-gene combinations (N=1,471/7,353) after adjusting for gene expression.

We hypothesized that peaks with a predicted allele-specific effect would be enriched for heritability of kidney-function-related traits in the proximal tubule. We partitioned heritability for eGFR using peaks that met the p-value threshold (Wald test p < 0.05) for the expression fixed effect in each of three models. All three models showed increased heritability for eGFR (Figure 7E). In contrast, proximal tubule ATAC peaks that showed increased proportion of fragments mapping to the alternate or reference allele (Figure 7C) in the aggregated dataset did not have enrichment for heritability of eGFR (ASCA, Figure 7D). These peaks do not necessarily have a predicted allele-specific effect and may represent a random subset of proximal tubule peaks that exhibit biased chromatin accessibility due to chance alone. These data suggest that peaks with a predicted allele-specific effect are more likely to contribute to heritability of eGFR than a random sampling of proximal tubule peaks. Approximately 20% of proximal tubule DAR were overlapping with a peak with a predicted allelic effect (N=104/597, Figure 7E, Hypergeometric p=1.7×10^-10), suggesting that genetic background may also modify chromatin accessibility patterns in DKD. We used the 4,476 significant peak-gene combinations present in all three models to perform gene ontology enrichment. The most enriched pathways involved peptide antigen assembly with MHC class II, antigen processing, and immunoglobulin production. Each of these pathways involve multiple HLA genes, which are known to exhibit allele-specific expression due to genetic variation in CRE (64). Proximal tubule expression of MHC class II regulates the response to kidney injury and renal fibrosis (65). Additional enriched pathways with important function in the proximal tubule included triglyceride metabolism, amino acid transport, and carbohydrate metabolism.

## Discussion

Single cell sequencing has advanced our understanding of kidney biology and multimodal analysis provides even greater insight into disease pathogenesis. DKD progression is multifactorial and contributing factors include hyperglycemia, hypertension, hypoxia, and inflammation (66). These factors exert their effect on different cell types throughout the nephron, which organizes a coordinated response to tissue injury. DKD was associated with an increased proportion of *VCAM1+* proximal tubule cells (PT_VCAM1) and infiltrating leukocytes in both snRNA-seq and snATAC-seq datasets. The PT_VCAM1 cell state emerges after proximal tubule injury and is associated with acute kidney injury, aging, and DKD (3,67). It adopts a pro-inflammatory phenotype characterized by enhanced NFκB signaling and failed repair that may underlie transition from acute kidney injury to CKD (68).

GR signaling is a key regulator of the immune response and exerts its effects on multiple cell types in the kidney. GR has potent anti-inflammatory properties that help mitigate tissue injury, but long-term exposure to glucocorticoids can lead to insulin resistance and metabolic syndrome (69). Chromatin accessibility pre-determines cellular response to glucocorticoids and GR preferentially binds areas of open chromatin (41). Our snATAC-seq analysis showed that the majority of DAR in DKD had reduced chromatin accessibility and were enriched for GR motifs across multiple cell types. These data suggest that the diabetic nephron is pre-programmed to respond differently to corticosteroids. Metabolic memory is an epigenetic state characterized by persistent expression of DKD-related genes despite glycemic control (5). Decreased chromatin accessibility of GR binding sites within GR-responsive genes can lead to reduced transactivation and expression of target genes, however, DNA-binding-independent mechanisms may remain intact. GR directly binds pro-inflammatory transcription factors like NFκB to inhibit their activity in a process called tethering (70). In a simple model, the metabolic effects of GR signaling can be attributed to transactivation and the anti-inflammatory effects can be attributed to tethering (71,72). We hypothesize that the diabetic kidney adapts to a pro-inflammatory environment by remodeling chromatin accessibility to promote anti-inflammatory effects of GR at the expense of its adverse effects on metabolism.

We used CUT&RUN to identify GR binding sites in the proximal tubule and validate predictions from our snATAC-seq analysis. GR binding sites showed significant overlap with proximal-tubule-specific ATAC peaks and participated in CCAN with cell-specific DAR in diabetes. A subset of GR CUT&RUN sites showed reduced chromatin accessibility in the proximal tubule, suggesting that it may respond differently to glucocorticoids. Changes in GR signaling were compounded by increased expression of GR, and reduced expression of *FKBP5* and *HSD11B2* in diabetes. *FKBP5* is a cytosolic chaperone that negatively regulates GR signaling as part of a negative feedback loop and *HSD11B2* converts cortisol into inactive cortisone to protect non-selective activation of MR (16). We found multiple DAR within *FKBP5* that coincide with GR binding sites within a CCAN. CRISPRi targeting of GR binding sites decreased *FKBP5* expression, suggesting that DKD is associated with reduced activity of GR negative feedback in the proximal tubule. *FKBP5* methylation has been associated with type 2 diabetes and cardiovascular risk and *FKBP5* polymorphisms are associated with insulin resistance (15,17). We hypothesize that *FKBP5* hypermethylation may lead to reduced chromatin accessibility in GRE and reduced activity of the GR negative feedback loop (Figure 8).

**Figure 8.**
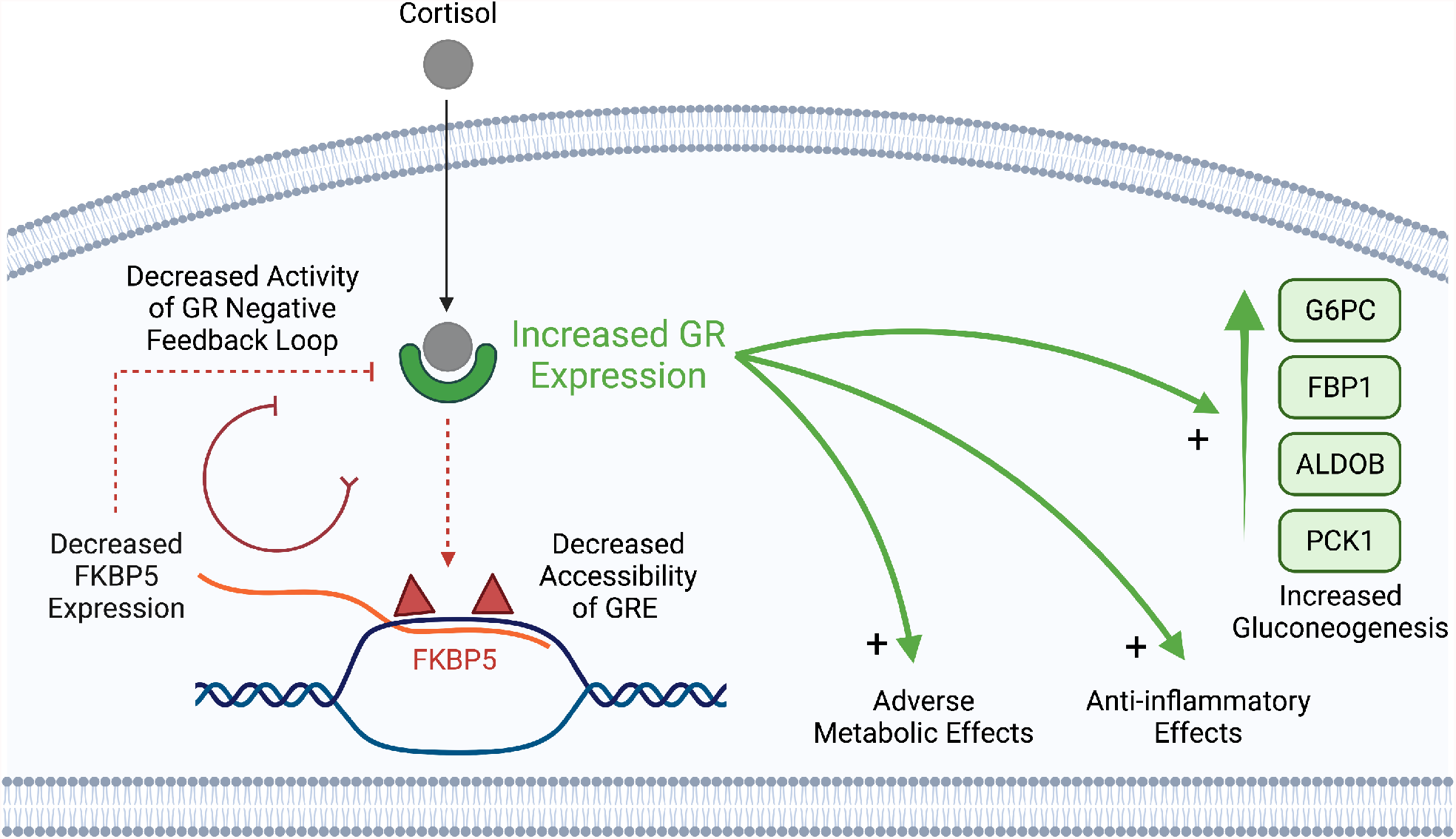
Model of altered glucocorticoid receptor signaling in the diabetic proximal tubule. GR expression is increased in the diabetic proximal tubule. Cortisol binds GR and translocates to the nucleus where it localizes to glucocorticoid response elements (GRE) in genes like *FKBP5*. Decreased chromatin accessibility of GRE in the *FKBP5* gene body is observed as reduced accessibility of proximal-tubule-specific ATAC peaks in DKD (red trianges). Reduced accessibility of *FKBP5* GRE leads to reduced transactivation by GR and reduced *FKBP5* expression. Reduced *FKBP5* expression decreases activity of the GR negative feedback loop. In the absence of *FKBP5* negative feedback, GR can exert both DNA-binding-dependent and DNA-binding-independent actions that lead to adverse metabolic effects, anti-inflammatory effects, and increased gluconeogenesis.

The diabetic proximal tubule had increased gluconeogenesis, which is downstream of GR signaling. The proximal tubule is the primary site in the kidney for glucose production and its rate-limiting enzyme is *PCK1* (6). We saw increased expression of *PCK1* and other gluconeogenic enzymes in the diabetic proximal tubule that was associated with reduced expression of *INSR*. Glucose reabsorption and glucose production are closely-intertwined and tightly regulated by insulin signaling (73). Proximal-tubule-specific *INSR* knockout and proximal-tubule-specific *IRS1/2* knockout have both been shown to increase gluconeogenesis, which is normally suppressed by insulin signaling or glucose reabsorption via SGLT2 (73,74). Glutamine is the preferred substrate for gluconeogenesis in the proximal tubule, which leads to ammonia production and acid excretion to maintain acid-base balance during prolonged fasting. It is possible that gluconeogenesis is upregulated in DKD to promote acid excretion and mitigate metabolic acidosis (75). SGLT2i stimulate gluconeogenesis in the liver and kidney, however, it remains unclear whether gluconeogenesis affects DKD progression (73,76,77).

Genetic background is increasingly recognized as an important determinant of kidney function and DKD (19,51,54). GWAS are generating a growing list of variants associated with kidney function, but it remains difficult to associate these variants with regulation of a specific gene or pathway. Bioinformatics approaches have led to the identification of quantitative trait loci associated with expression (eQTL), chromatin accessibility (caQTL), methylation (meQTL), and other traits that may regulate kidney function in a cell-specific manner (23). Some of these phenotypes are driven by allele-specific effects that can be measured as changes in expression (ASE) and chromatin accessibility (ASCA) (3,29,78). Allele-specific analysis can substantially boost the power of QTL studies because each individual serves as its own perfectly-matched control. Multimodal single cell datasets can take advantage of this approach to model allele-specific effects as a function of gene expression (or any other measurable quantity). We developed a tool for single cell allele-specific analysis called SALSA and used it to detect proximal-tubule-specific ATAC peaks in CRE that modify gene expression via ASCA. These peaks were enriched for heritability of eGFR and some of them coincide with DAR in DKD. These findings raise the possibility that genetic background affects kidney function via ASCA, which could alter progression of DKD.

## Methods

### Human Kidney Tissue

Non-tumor kidney cortex samples (n=10) were obtained from patients undergoing partial or radical nephrectomy for renal mass at Brigham and Women’s Hospital (Boston, MA) under an established Institutional Review Board protocol approved by the Mass General Brigham Human Research Committee. An additional three kidney cortex samples (1 control and 2 DKD) were obtained from deceased organ donors in the Novo Nordisk biorepository. All participants provided written informed consent in accordance with the Declaration of Helsinki. Histologic sections were reviewed by a renal pathologist and laboratory data was abstracted from the medical record.

### Nuclear Dissociation and Library Preparation

Samples were chopped into < 2 mm pieces, homogenized with a Dounce homogenizer (885302–0002; Kimble Chase) in 2 ml of ice-cold Nuclei EZ Lysis buffer (NUC-101; Sigma-Aldrich) supplemented with protease inhibitor (5892791001; Roche) with or without RNase inhibitors (Promega, N2615 and Life Technologies, AM2696, only for snRNA-seq library preparation), and incubated on ice for 5 min. The homogenate was filtered through a 40-μm cell strainer (43–50040–51; pluriSelect) and centrifuged at 500g for 5 min at 4°C. The pellet was resuspended, washed with 4 ml of buffer, and incubated on ice for 5 min. Following centrifugation, the pellet was resuspended in Nuclei Buffer (10× Genomics, PN-2000153) for snATAC-seq, nuclei suspension buffer (1x PBS, 1% bovine serum albumin [BSA], 0.1% RNase inhibitor) for snRNA-seq, or 1xPBS containing 1% BSA for CUT&RUN. The suspension was then filtered through a 5-μm cell strainer (43-50005-03, pluriSelect) and counted.

### Single Nucleus ATAC Sequencing and Bioinformatics Workflow

Thirteen snATAC-seq libraries were created with 10X Genomics Chromium Single Cell ATAC v1 chemistry following nuclear dissociation. These libraries included six healthy control and seven DKD samples. Five of the healthy control snATAC-seq libraries were prepared for a prior study (GSE151302). A target of 10,000 nuclei were loaded onto each lane. Sample index PCR was performed at 12 cycles. Libraries were sequenced on an Illumina Novaseq instrument and counted with cellranger-atac v2.0 (10X Genomics) using GRCh38. Libraries were aggregated with cellranger-atac without depth normalization. A mean of 327,328,680 reads were sequenced for each snATAC library (s.d. = 47,171,305) corresponding to a median of 15,150 fragments per cell (s.d. = 3,875). The mean fraction of reads with a valid barcode was 96.3 ± 2.2%. Aggregated datasets were processed with Seurat v4.0.3 and its companion package Signac v1.3.0. Low-quality cells were removed from the aggregated snATAC-seq dataset (peak region fragments > 2500, peak region fragments < 20000, nucleosome signal < 4, TSS enrichment > 2) before normalization with term-frequency inverse-document-frequency (TFIDF). Doublets were removed with AMULET (79). Dimensional reduction was performed via singular value decomposition (SVD) of the TFIDF matrix followed by UMAP. A KNN graph was constructed to cluster cells with the Louvain algorithm. Batch effect was corrected with Harmony using the RunHarmony function in Seurat. A gene activity matrix was constructed by counting ATAC peaks within the gene body and 2 kb upstream of the transcriptional start site using protein-coding genes annotated in the Ensembl database. The gene activity matrix was log-normalized prior to label transfer with the aggregated snRNA-seq Seurat object using canonical correlation analysis. The aggregated snATAC-seq object was filtered using label transfer to remove additional heterotypic doublets not captured by AMULET. Cell-specific ATAC peaks were called with MACS2 using the Signac wrapper and a new Seurat object was created using MACS2 peaks and the FeatureMatrix function. The new snATAC-seq object was reprocessed with TFIDF, SVD, and batch effect correction followed by clustering and annotation based on lineage-specific gene activity.

After filtering, there was a mean of 6000 ± 1134 nuclei per snATAC-seq library with a mean of 8098 ± 3231 peaks detected per nucleus. The final snATAC-seq library contained a total of 437,311 unique peak regions among 68,458 nuclei and represented all major cell types within the kidney cortex. Differential chromatin accessibility between cell types was assessed with the Signac FindMarkers function for peaks detected in at least 20% of cells using a likelihood ratio test. Bonferroni-adjusted p-values were used to determine significance at an FDR < 0.05. Genomic regions containing snATAC-seq peaks were annotated with ChIPSeeker (v1.5.1) and clusterProfiler (v4.0.5) using Ensembl and FANTOM databases on hg38. Motif enrichment within DAR was calculated with the Signac FindMotifs function using cell-specific accessible peaks matched for GC content. chromVAR motif activities were computed using the Signac wrapper and JASPAR2020 database adjusted for the number of fragments in peaks for each nucleus. CCAN were computed with cicero (v1.3.4.11) using the run_cicero function and default parameters. Gene-enhancer links were computed with the Signac LinkPeaks function and imputed RNA following label transfer and integration of snRNA-seq and snATAC-seq datasets.

### Single Nucleus RNA Sequencing and Bioinformatics Workflow

Eleven snRNA-seq libraries were obtained using 10X Genomics Chromium single cell chemistry following nuclear dissociation. Eight snRNA-seq libraries (5 control, 3 DKD) were prepared for prior studies (GSE13188213, GSE151302). A target of 10,000 nuclei were loaded onto each lane. The cDNA for snRNA libraries was amplified for 17 cycles. Libraries were sequenced on an Illumina Novaseq instrument and counted with cellranger v4.0 using a custom pre-mRNA GTF built on GRCh38 to include intronic reads. Datasets were aggregated with cellranger v4.0 without depth normalization. A mean of 382,207,065 reads (s.d. = 78,522,614) were sequenced for each snRNA library corresponding to a mean of 68,429 reads per cell (s.d. = 21,706). The mean sequencing saturation was 77.6 ± 11.9%. The mean fraction of reads with a valid barcode (fraction of reads in cells) was 75.9 ± 6.6%. Aggregated datasets were preprocessed with Seurat v4.0.3 to remove low-quality nuclei (Features > 500, Features < 5000, RNA count < 16000, %Mitochondrial genes < 0.5, %Ribosomal protein large or small subunits < 0.3) and DoubletFinder v2.0.3 to remove heterotypic doublets (assuming 6% of barcodes represent doublets).

The filtered library was normalized with SCTransform, and corrected for batch effects with Harmony v0.1.0 using the RunHarmony function in Seurat. After filtering, there was a mean of 3561 ± 2028 cells per snRNA-seq library and a mean of 2137 ± 1031 genes detected per nucleus. Clustering was performed by constructing a KNN graph and applying the Louvain algorithm. Dimensional reduction was performed with UMAP and individual clusters were annotated based on expression of lineage-specific markers. The final snRNA-seq library contained 39,176 cells and represented all major cell types within the kidney cortex. Differential expression between cell types was assessed with the Seurat FindMarkers function for transcripts detected in at least 20% of cells. Bonferroni-adjusted p-values were used to determine significance at an FDR < 0.05.

### Single Cell Analysis of Five-prime Ends with SCAFE

We used SCAFE to analyze single nucleus 5’ paired-end chemistry libraries obtained from two DKD samples and two previously-published healthy control samples (GSE151302) (36). Transcribed cis-regulatory elements (tCRE) were called in individual libraries using the scafe.worfklow.sc.solo function with default parameters. There was a mean of 16,912 tCRE per library (s.d. = 1855). Libraries were pooled with the scafe.workflow.sc.pool function to identify a total of 37,698 tCRE. Pooled libraries were merged into a Seurat object followed by normalization with SCTransform, batch effect correction with Harmony, dimensional reduction, and clustering. The final object contained 10,984 nuclei and cell types were annotated using snRNA-seq barcode annotations from the same samples (see snRNA-seq bioinformatics workflow above). Cell-specific tCRE and differential tCRE in diabetes were identified with the FindMarkers function with a log-fold-change threshold of 0.25. Bonferroni adjusted p-values were used to determine significance at an FDR < 0.05 and significant tCRE were annotated with ChIPseeker.

### Cell Culture

Human primary proximal tubular cells (human RPTEC, Lonza; CC-2553) were cultured with renal epithelial cell growth medium kit (Lonza; CC-3190). Human telomerase reverse transcriptase (hTERT)-immortalized human RPTEC (ATCC; CRL-4031) were cultured with ATCC hTERT Immortalized RPTEC Growth Kit (ATCC, ACS-4007). HEK293T cells (ATCC; CRL-3216) were cultured in Dulbecco’s modified Eagle’s medium (DMEM, Gibco; 11965092) supplemented with 10% fetal bovine serum (Gibco; 10437028) and antibiotics. All cultured cells were maintained in a humidified 5% CO2 atmosphere at 37°C.

### GR CUT&RUN Library Preparation and Peak Calling

CUT&RUN assay libraries for cultured cells or human kidneys were generated with the CUTANA kit (EpiCypher, 14-1048). For cultured cells, adherent cells were scraped from culture dishes and centrifuged at 500 × g for 5 min. Pellets were resuspended in PBS with 1% BSA and counted. Cultured cells or nuclei obtained from a human kidney (500,000 cells or nuclei) were then mixed and incubated with Concanavalin A (ConA) conjugated paramagnetic beads. Antibodies were added to each sample (0.5μg of rabbit glucocorticoid receptor antibody [abcam, ab225886, 1:20] or rabbit IgG negative control antibody [Epicypher, 13-0041k, 1:50]). The remaining steps were performed according to manufacturer’s instructions. Library preparation was performed using the NEBNext Ultra II DNA Library Prep Kit for Illumina (New England BioLabs, E7645S) with manufacturer’s instructions, including minor modifications indicated by CUTANA described above. CUT&RUN libraries were sequenced on a NovaSeq instrument (Illumina, 150 bp paired-end reads). Fastq files were trimmed with Trim Galore (Cutadapt [v2.8]) and aligned with Bowtie2 [v2.3.5.1] (parameters: --local --very-sensitive-local --no-unal --no-mixed --no-discordant --phred33 -I 10 -X 700) using hg38. Peak calling was performed using MACS2 [v2.2.7.1] with default parameters.

### Bulk ATAC-seq Library Preparation and Peak Calling

We suspended 50,000 cells in 50 μL of ice-cold lysis buffer with 10 mM Tris-HCl (pH 7.4), 10 mM NaCl, 3 mM MgCl2, 1% BSA, 0.1% Tween-20 (Sigma, P7949-100ML), 0.1% NP-40 (Thermo Scientific, 28324) and 0.01% Digitonin (Thermo Scientific, BN2006) (80). The suspension was incubated for 4 min on ice. Subsequently, 450 μL of ice-cold wash buffer (10 mM Tris-HCl [pH 7.4], 10 mM NaCl, 3 mM MgCl2, 1% BSA, 0.1% Tween-20) was added and centrifuged at 600 × g for 6 min. The pellet was resuspended in 25 μL of ATAC-seq transposition mix (12.5 μL 2× Illumina Tagment DNA (TD) buffer; 10.5 μL nuclease-free water; 2.0 μL Tn5 transposase [Illumina, FC-121-1030]) and incubated at 37°C for 1 h on a thermomixer. The transposed DNA was purified with MinElute PCR purification kit (QUIAGEN, 28004). DNA samples were then amplified with PCR ([72°C; 5 min] and [98°C; 30 s] followed by 9 cycles of [98°C; 10 s, 63°C; 30 s, 72°C; 1 min] using unique 10-bp dual indexes and NEBNext High-Fidelity 2× PCR Master Kit (M0541L).

Following the first amplification, DNA size selection was performed using solid-phase reversible immobilization (SPRI) beads (AMPure XP [Beckman Coulter, A63881]) at an SPRI to DNA ratio of 0.5. The supernatant was further mixed with SPRI beads at a SPRI to DNA ratio of 1.2. The resulting supernatant was discarded, and the magnet-immobilized SPRI beads were washed twice with 80% ethanol. DNA was subsequently eluted in 20 μL of EB elution buffer (QUIAGEN, included in 28004). The size-selected DNA was amplified with an additional 9-cycle PCR. Subsequently, the amplified DNA was purified with Ampure XP (SPRI to DNA ratio of 1.7) and eluted with 25 μL of buffer EB elution buffer. The resultant ATAC-seq libraries were sequenced on a NovaSeq instrument (Illumina, 150 bp paired-end reads). Fastq files were trimmed with Trim Galore (Cutadapt [v2.8]) and aligned with Bowtie2 [v2.3.5.1] with --very-sensitive -X 2000 using hg38. PCR duplicates were removed with Picard’s MarkDuplicates function. Peak calling was performed on each sample separately using MACS2 [v2.2.7.1] (--nomodel --shift -100 -- extsize 200). The consensus list of accessible peaks was generated using the intersectBed function in bedtools.

### CRISPR Interference

Small guide RNA (sgRNA) targeting around the *FKBP5* TSS and intronic CRE were designed with CHOPCHOP (https://chopchop.cbu.uib.no/). These sgRNAs and two non-targeting control sgRNAs were placed following the U6 promoter in a dCas9-KRAB repression plasmid (pLV hU6-sgRNA hUbC-dCas9-KRAB-T2a-Puro, Addgene; 71236, a gift from Charles Gersbach) with golden gate assembly. The sgRNA sequences used in this study are in Supplemental Table 20. First, single-strand oligonucleotides (Integrated and Technology [IDT]) for sense and anti-sense sequences were annealed. Subsequently, cloning with Golden gate assembly was performed with Esp3I restriction enzyme (NEB, R0734L) and T4 DNA ligase (NEB, M0202L) on a thermal cycler repeating 37°C for 5 min and 16°C for 5 min for 60 cycles, followed by transformation to NEB 5-alpha Competent E. coli (NEB, C2987H) per manufacturer’s instructions. The cloned lentiviral vectors were purified with a mini high-speed plasmid kit (IBI Scientific; IB47102). Insertion of sgRNA was checked with Sanger sequencing. For lentivirus preparation, we seeded 6.0×10^5^ HEK293T cells per well on 6-well tissue culture plates 16 h prior to transfection. Cells were transfected with 1.5 µg of psPAX2 (Addgene; 12260, a gift from Didier Trono), 0.15 µg of pMD2.G (Addgene; 12259, a gift from Didier Trono) and 1.5 µg of dCas9-KRAB repression plasmid per well by Lipofectamine 3000 transfection reagent (Invitrogen; L3000015) per manufacturer’s instructions.

Culture media were changed to DMEM supplemented with 30% FBS 24 h after transfection. Lentivirus-containing supernatants were harvested 24 h later and filtered with 0.45 µm PVDF filters (CELLTREAT; 229745). The lentivirus-containing supernatants were immediately used for lentiviral transduction. Human RPTEC were seeded at 5.0×10^4^ cells per well on 6-well tissue culture plates 16 h prior to transfection. The media on human RPTEC was then changed to the fresh lentiviral supernatants supplemented with polybrene (0.5 µg/ml, Santa Cruz Biotechnology; sc-134220) and cultured for 24 h. Subsequently, RPTEC cells were cultured in DMEM with 10% FBS and puromycin (3 µg/ml, invivogen; ant-pr-1) for 72 h.

### Quantitative PCR

RNA from human RPTECs was extracted with the TRIZOL and Direct-zol MicroPrep Plus Kit (Zymo) following manufacturer’s instructions. Extracted RNA (1-2 µg) was used for reverse transcription to generate cDNA libraries with the High-Capacity cDNA Reverse Transcription Kit (Life Technologies). Quantitative PCR was performed in the BioRad CFX96 Real-Time System using iTaq Universal SYBR Green Supermix (Bio-Rad). Expression levels were normalized to *GAPDH*, and data were analyzed using the 2-ΔΔCt method. Quantitative PCR data are presented as mean±s.d. and compared between groups with one-way ANOVA and a post-hoc Dunnett’s adjustment for multiple comparisons. A p-value < 0.05 was considered statistically significant. Primer sequences are provided in supplementary materials (Supplemental Table 20).

### Bulk RNA-seq Analysis of Previously-published Human DKD

Raw fastq files were downloaded from GSE142025 to include 9 healthy control, 6 early DKD, and 22 advanced DKD donors (49). Transcript abundance was quantified with Salmon using Ensembl (release-99) and count matrices were imported to DESeq2 with tximport (v1.16.1). Differentially expressed genes were identified using the DESeq function with default parameters for early DKD vs. Control and advanced DKD vs. Control (Supplemental Table 18). Significance was determined using a Benjamini-Hochberg adjusted p-value.

### Partitioned Heritability of ATAC Peaks

Cell-specific ATAC peaks and cell-specific DAR in diabetes were identified with the Seurat FindMarkers function, sorted by p-value, and filtered for peaks with an average log-fold-change greater than zero. All peaks that met the adjusted p-value threshold were used to generate a cell-specific bed file. In the event a cell type did not have at least 2000 peaks that met the adjusted p-value threshold, the top 2000 peaks with the lowest p-value were used to create the bed file. If a cell type did not have 2000 cell-specific peaks, all available peaks were used to create the bed file. For the allele-specific analysis, bed files were generated using peaks that met the adjusted binomial threshold for allelic bias of reference vs. alternate allele (N=5593, padj < 0.05) or the unadjusted p-value threshold for the base model (N=7512, pval < 0.05), model 2 (N=7557, pval < 0.05), and model 3 (N=7353, pval < 0.05). These thresholds were used to keep the number of peaks in each annotation roughly equivalent. Bed files were lifted over to hg19 to create annotations for autosomal chromosomes with a 1000 genomes phase 3 reference and the make_annot.py function in ldsc using a 100kb window (55). Linkage disequilibrium scores were computed from custom annotations with the ldsc.py function using default parameters. GWAS summary statistics for eGFR, CKD, microalbuminuria, and urinary sodium excretion were downloaded from publicly-available databases and formatted for ldsc using munge_sumstats.py (52–54). Partitioned heritability for each GWAS trait was estimated using the 1000G phase 3 reference and ldsc cell-type-specific workflow with default parameters, including baseline v1.2 annotations after controlling for all kidney ATAC peaks in the dataset. P-values were adjusted for multiple comparisons using Benjamini-Hochberg and significance was determined at padj < 0.05.

### Allele-specific Modeling with SALSA

Coordinate-sorted bam files generated by cellranger (snRNA-seq) or cellranger-atac (snATAC-seq) were genotyped with SALSA using GATK best practices for germline short variant discovery (58). For snRNA-seq, reads containing Ns in their cigar string (eg. spanning splice junctions in snRNA-seq data) were split using SplitNCigarReads. For snRNA-seq and snATAC-seq, base recalibration was performed with BaseRecalibrator using hg38 GATK bundle resources, including dbsnp (v138), 1000G phase I indels, 1000G phase I high-confidence SNV, and Mills and 1000G gold standard indels. Recalibration was applied with ApplyBQSR to create analysis-ready bam files. Variants were identified from analysis-ready bam files with HaplotypeCaller and genotypes were called from GVCFs using GenotypeGVCFs with default parameters. snRNA-seq variants were hard-filtered by Fisher strand bias (FS > 30), quality by depth score (QD < 2), cluster size (3), and cluster-window size (35bp). snATAC-seq were filtered using CNNScoreVariants followed by FilterVariantTranches (--snp-tranche 99.95 indel-tranche 99.4). snRNA-seq libraries had a mean total of 98,255 (s.d. = 64,847) SNVs and indels and snATAC-seq libraries had a mean total of 2,283,904 (s.d. = 534,165) SNVs and indels. Genotypes from snRNA-seq and snATAC-seq were combined and phased with shapeit4.2 using the 1000 Genomes phased reference for biallelic SNV and indels on GRCh38 (58). There was a mean of 1,917,939 (s.d. = 363,223) phased SNV and indels for each library, which were used to perform variant-aware realignment with WASP (60). WASP-aligned bam files were divided into single cell bam files by extracting proximal-tubule-specific barcodes using the CB tag. GATK ASEReadCounter was used to generate single cell allele-specific counts from single cell bam files using phased heterozygous SNV (mean = 722,091, s.d. = 199,756).

Pseudo-multiomic cells were created by performing label transfer from the aggregated snRNA-seq to snATAC-seq dataset to generate an imputed RNA estimate for each snATAC-seq cell. One or more gene targets for each ATAC peak were identified using the LinkPeaks function. ATAC peaks with heterozygous SNV were filtered for an aggregate total fragment count greater than 20, total reference allele count greater than 5, and total alternate allele count greater than 5. The ratio of aggregated reference counts to alternate counts within ATAC peaks containing heterozygous SNV was compared with a binomial test. A generalized linear mixed effect model with a logit link function was implemented with the lme4 package (81). In the base model, the dependent variable was coded as the presence of an alternate allele within an ATAC peak and the continuous predictor variable was the imputed RNA estimate normalized to an interval from 0 to 100. A mixed effect per sample was added to control for pseudo-replication bias. In model 2, an additional fixed effect for diabetes was added. In model 3, additional fixed effects for diabetes and an interaction term between diabetes and imputed RNA expression were added. Significance of each peak-gene combination was evaluated using a Wald test obtained from the glmer function. In the supplementary materials, all fixed-effect coefficient estimates for each peak-gene combination are included with 95% confidence intervals, Wald p-values, and standard deviation estimates of random effects (Supplemental Table 19). Peak-gene combinations meeting the p-value threshold for expression (p < 0.05) were annotated with ChIPseeker and the corresponding effect size in log-odds is visualized in relation to the nearest TSS. These same peaks were also used to create annotations and calculate linkage disequilibrium scores with ldsc for partitioned heritability of eGFR as previously described.

## Data Availability

All relevant data are available from the corresponding authors on reasonable request. Raw sequencing data for snATAC-seq (N=1 control, N=7 DKD) and snRNA-seq (N=1 control, N=2 DKD) is deposited in GEO under accession number GSEXXXXXXX (Reviewer token: XXXXXXXXXXXX). Previously published raw sequencing data for snRNA-seq (N=5 control, N=3 DKD) and snATAC-seq (N=5 control) are available in GEO (GSE151302, GSE131882). Processed count matrices for all snRNA-seq (N=11) and snATAC-seq (N=13) libraries used in this study are provided in GSEXXXXXXX. Sequencing data for CUT&RUN from bulk kidney cortex and primary RPTEC are deposited under accession number GSEXXXXX (Reviewer token: XXXXXXXXX). Sequencing data for Omni-ATAC from hTERT-RPTEC and primary RPTEC are also deposited under accession number GSEXXXXXXXX. Gene expression and chromatin accessibility for each cell type can be viewed on our interactive website; Kidney Interactive Transcriptomics (http://humphreyslab.com/SingleCell) (Reviewer password: XXXXXXXXX).

## Code Availability

SALSA is available on GitHub (https://github.com/p4rkerw/SALSA). All code used to generate the data in this manuscript will also be made available on GitHub (https://github.com/p4rkerw) when this manuscript completes peer review.

## Acknowledgements

These studies were supported by a pilot grant from the Washington University Diabetes Research Center NIH DK20579 to P.C.W., NIH K08 Career Development Award DK126847 to P.C.W, DK103740 to B.D.H. and Chan Zuckerberg Initiative seed network grant CZF2019-002430 to B.D.H. Additional support was provided by the International Research Fund for Subsidy of Kyushu University School of Medicine Alumni, the Japan Society for the Promotion of Science (JSPS) Postdoctoral Fellowships for Research Abroad, and the Osamu Hayaishi Memorial Scholarship for Study Abroad to Y.M.

## Literature Cited

1. Johansen KL, Chertow GM, Foley RN, Gilbertson DT, Herzog CA, Ishani A, et al. US Renal Data System 2020 Annual Data Report: Epidemiology of Kidney Disease in the United States. American Journal of Kidney Diseases. 2021 Apr 1;77(4, Supplement 1):A7–8.

2. Boer IH de, Caramori ML, Chan JCN, Heerspink HJL, Hurst C, Khunti K, et al. KDIGO 2020 Clinical Practice Guideline for Diabetes Management in Chronic Kidney Disease. Kidney International. 2020 Oct 1;98(4):S1–115.

3. Muto Y, Wilson PC, Ledru N, Wu H, Dimke H, Waikar SS, et al. Single cell transcriptional and chromatin accessibility profiling redefine cellular heterogeneity in the adult human kidney. Nat Commun. 2021 Apr 13;12(1):2190.

4. Stuart T, Butler A, Hoffman P, Hafemeister C, Papalexi E, Mauck WM, et al. Comprehensive Integration of Single-Cell Data. Cell. 2019 Jun 13;177(7):1888-1902.e21.

5. Kato M, Natarajan R. Epigenetics and epigenomics in diabetic kidney disease and metabolic memory. Nat Rev Nephrol. 2019 Jun;15(6):327–45.

6. Legouis D, Faivre A, Cippà PE, de Seigneux S. Renal gluconeogenesis: an underestimated role of the kidney in systemic glucose metabolism. Nephrology Dialysis Transplantation [Internet]. 2020 Nov 28 [cited 2021 Jun 17];(gfaa302). Available from: https://doi.org/10.1093/ndt/gfaa302

7. Kang A, Jardine MJ. SGLT2 inhibitors may offer benefit beyond diabetes. Nat Rev Nephrol. 2021 Feb;17(2):83–4.

8. Wilson PC, Wu H, Kirita Y, Uchimura K, Ledru N, Rennke HG, et al. The single-cell transcriptomic landscape of early human diabetic nephropathy. PNAS. 2019 Sep 24;116(39):19619–25.

9. Bauerle KT, Harris C. Glucocorticoids and Diabetes. Mo Med. 2016;113(5):378–83.

10. Hwang JL, Weiss RE. Steroid-induced diabetes: a clinical and molecular approach to understanding and treatment. Diabetes Metab Res Rev. 2014 Feb;30(2):96–102.

11. Gerich JE, Meyer C, Woerle HJ, Stumvoll M. Renal Gluconeogenesis: Its importance in human glucose homeostasis. Diabetes Care. 2001 Feb 1;24(2):382–91.

12. Khani S, Tayek JA. Cortisol increases gluconeogenesis in humans: its role in the metabolic syndrome. Clin Sci (Lond). 2001 Dec;101(6):739–47.

13. Hartmann J, Bajaj T, Klengel C, Chatzinakos C, Ebert T, Dedic N, et al. Mineralocorticoid receptors dampen glucocorticoid receptor sensitivity to stress via regulation of FKBP5. Cell Rep. 2021 Jun 1;35(9):109185.

14. Hubler TR, Scammell JG. Intronic hormone response elements mediate regulation of FKBP5 by progestins and glucocorticoids. Cell Stress Chaperones. 2004 Jul;9(3):243–52.

15. Ortiz R, Joseph JJ, Lee R, Wand GS, Golden SH. Type 2 diabetes and cardiometabolic risk may be associated with increase in DNA methylation of FKBP5. Clinical Epigenetics. 2018 Jun 19;10(1):82.

16. Zannas AS, Wiechmann T, Gassen NC, Binder EB. Gene–Stress–Epigenetic Regulation of FKBP5 : Clinical and Translational Implications. Neuropsychopharmacol. 2016 Jan;41(1):261–74.

17. Fichna M, Krzyśko-Pieczka I, Zurawek M, Skowronska B, Januszkiewicz-Lewandowska D, Fichna P. FKBP5 polymorphism is associated with insulin resistance in children and adolescents with obesity. Obes Res Clin Pract. 2018 Feb;12(Suppl 2):62–70.

18. Zhang Y, Liu T, Meyer CA, Eeckhoute J, Johnson DS, Bernstein BE, et al. Model-based analysis of ChIP-Seq (MACS). Genome Biol. 2008;9(9):R137.

19. van Zuydam NR, Ahlqvist E, Sandholm N, Deshmukh H, Rayner NW, Abdalla M, et al. A Genome-Wide Association Study of Diabetic Kidney Disease in Subjects With Type 2 Diabetes. Diabetes. 2018;67(7):1414–27.

20. Guan M, Keaton JM, Dimitrov L, Hicks PJ, Xu J, Palmer ND, et al. Genome-wide association study identifies novel loci for type 2 diabetes-attributed end-stage kidney disease in African Americans. Human Genomics. 2019 May 15;13(1):21.

21. Xue A, Wu Y, Zhu Z, Zhang F, Kemper KE, Zheng Z, et al. Genome-wide association analyses identify 143 risk variants and putative regulatory mechanisms for type 2 diabetes. Nat Commun. 2018 Jul 27;9(1):1–14.

22. Doke T, Huang S, Qiu C, Liu H, Guan Y, Hu H, et al. Transcriptome-wide association analysis identifies DACH1 as a kidney disease risk gene that contributes to fibrosis. J Clin Invest. 2021 May 17;131(10):141801.

23. Sullivan KM, Susztak K. Unravelling the complex genetics of common kidney diseases: from variants to mechanisms. Nat Rev Nephrol. 2020 Nov;16(11):628–40.

24. Rai V, Quang DX, Erdos MR, Cusanovich DA, Daza RM, Narisu N, et al. Single-cell ATAC-Seq in human pancreatic islets and deep learning upscaling of rare cells reveals cell-specific type 2 diabetes regulatory signatures. Molecular Metabolism. 2020;32:109–21.

25. Chiou J, Geusz RJ, Okino M-L, Han JY, Miller M, Melton R, et al. Interpreting type 1 diabetes risk with genetics and single-cell epigenomics. Nature. 2021 Jun;594(7863):398–402.

26. Minnoye L, Marinov GK, Krausgruber T, Pan L, Marand AP, Secchia S, et al. Chromatin accessibility profiling methods. Nat Rev Methods Primers. 2021 Jan 21;1(1):1–24.

27. Liang D, Elwell AL, Aygün N, Krupa O, Wolter JM, Kyere FA, et al. Cell-type-specific effects of genetic variation on chromatin accessibility during human neuronal differentiation. Nat Neurosci. 2021 May 20;1–13.

28. Pliner HA, Packer JS, McFaline-Figueroa JL, Cusanovich DA, Daza RM, Aghamirzaie D, et al. Cicero Predicts cis-Regulatory DNA Interactions from Single-Cell Chromatin Accessibility Data. Mol Cell. 2018 06;71(5):858-871.e8.

29. Atak ZK, Taskiran II, Demeulemeester J, Flerin C, Mauduit D, Minnoye L, et al. Interpretation of allele-specific chromatin accessibility using cell state–aware deep learning. Genome Res. 2021 Jun;31(6):1082–96.

30. Fang R, Preissl S, Li Y, Hou X, Lucero J, Wang X, et al. Comprehensive analysis of single cell ATAC-seq data with SnapATAC. Nat Commun. 2021 Feb 26;12(1):1337.

31. Korsunsky I, Millard N, Fan J, Slowikowski K, Zhang F, Wei K, et al. Fast, sensitive and accurate integration of single-cell data with Harmony. Nature Methods. 2019 Dec;16(12):1289–96.

32. Andersson R, Gebhard C, Miguel-Escalada I, Hoof I, Bornholdt J, Boyd M, et al. An atlas of active enhancers across human cell types and tissues. Nature. 2014 Mar 27;507(7493):455–61.

33. Spoto B, Pisano A, Zoccali C. Insulin resistance in chronic kidney disease: a systematic review. Am J Physiol Renal Physiol. 2016 Dec 1;311(6):F1087–108.

34. Moore JE, Purcaro MJ, Pratt HE, Epstein CB, Shoresh N, Adrian J, et al. Expanded encyclopaedias of DNA elements in the human and mouse genomes. Nature. 2020 Jul;583(7818):699–710.

35. Hirabayashi S, Bhagat S, Matsuki Y, Takegami Y, Uehata T, Kanemaru A, et al. NET-CAGE characterizes the dynamics and topology of human transcribed cis-regulatory elements. Nat Genet. 2019 Sep;51(9):1369–79.

36. Moody J, Kouno T, Suzuki A, Shibayama Y, Terao C, Chang J-C, et al. Profiling of transcribed cis-regulatory elements in single cells [Internet]. 2021 Apr [cited 2021 Dec 22] p. 2021.04.04.438388. Available from: https://www.biorxiv.org/content/10.1101/2021.04.04.438388v1

37. Kouno T, Moody J, Kwon AT-J, Shibayama Y, Kato S, Huang Y, et al. C1 CAGE detects transcription start sites and enhancer activity at single-cell resolution. Nat Commun. 2019 Jan 21;10(1):360.

38. Sartorelli V, Lauberth SM. Enhancer RNAs are an important regulatory layer of the epigenome. Nat Struct Mol Biol. 2020 Jun;27(6):521–8.

39. Tu Z, Kelley VR, Collins T, Lee FS. IκB Kinase Is Critical for TNF-α-Induced VCAM1 Gene Expression in Renal Tubular Epithelial Cells. J Immunol. 2001 Jun 1;166(11):6839–46.

40. Biddie SC, John S, Sabo PJ, Thurman RE, Johnson TA, Schiltz RL, et al. Transcription Factor AP1 Potentiates Chromatin Accessibility and Glucocorticoid Receptor Binding. Mol Cell. 2011 Jul 8;43(1):145–55.

41. John S, Sabo PJ, Thurman RE, Sung M-H, Biddie SC, Johnson TA, et al. Chromatin accessibility pre-determines glucocorticoid receptor binding patterns. Nat Genet. 2011 Mar;43(3):264–8.

42. Skene PJ, Henikoff S. An efficient targeted nuclease strategy for high-resolution mapping of DNA binding sites. Reinberg D, editor. eLife. 2017 Jan 12;6:e21856.

43. Khan A, Fornes O, Stigliani A, Gheorghe M, Castro-Mondragon JA, van der Lee R, et al. JASPAR 2018: update of the open-access database of transcription factor binding profiles and its web framework. Nucleic Acids Res. 2018 04;46(D1):D260–6.

44. Salyer SA, Parks J, Barati MT, Lederer ED, Clark BJ, Klein JD, et al. Aldosterone regulates Na+, K+ ATPase activity in human renal proximal tubule cells through mineralocorticoid receptor. Biochimica et Biophysica Acta (BBA) - Molecular Cell Research. 2013 Oct 1;1833(10):2143–52.

45. Funder JW. Reconsidering the Roles of the Mineralocorticoid Receptor. Hypertension. 2009 Feb 1;53(2):286–90.

46. Nan X, Ng HH, Johnson CA, Laherty CD, Turner BM, Eisenman RN, et al. Transcriptional repression by the methyl-CpG-binding protein MeCP2 involves a histone deacetylase complex. Nature. 1998 May 28;393(6683):386–9.

47. Schödel J, Ratcliffe PJ. Mechanisms of hypoxia signalling: new implications for nephrology. Nat Rev Nephrol. 2019 Oct;15(10):641–59.

48. Schep AN, Wu B, Buenrostro JD, Greenleaf WJ. chromVAR: inferring transcription-factor-associated accessibility from single-cell epigenomic data. Nat Methods. 2017 Oct;14(10):975–8.

49. Fan Y, Yi Z, D’Agati VD, Sun Z, Zhong F, Zhang W, et al. Comparison of Kidney Transcriptomic Profiles of Early and Advanced Diabetic Nephropathy Reveals Potential New Mechanisms for Disease Progression. Diabetes. 2019 Dec;68(12):2301–14.

50. Alerasool N, Segal D, Lee H, Taipale M. An efficient KRAB domain for CRISPRi applications in human cells. Nat Methods. 2020 Nov;17(11):1093–6.

51. Hellwege JN, Velez Edwards DR, Giri A, Qiu C, Park J, Torstenson ES, et al. Mapping eGFR loci to the renal transcriptome and phenome in the VA Million Veteran Program. Nat Commun. 2019 Aug 26;10(1):3842.

52. Wuttke M, Li Y, Li M, Sieber KB, Feitosa MF, Gorski M, et al. A catalog of genetic loci associated with kidney function from analyses of a million individuals. Nat Genet. 2019 Jun;51(6):957–72.

53. Teumer A, Li Y, Ghasemi S, Prins BP, Wuttke M, Hermle T, et al. Genome-wide association meta-analyses and fine-mapping elucidate pathways influencing albuminuria. Nat Commun. 2019 Sep 11;10(1):4130.

54. Pazoki R, Evangelou E, Mosen-Ansorena D, Pinto RC, Karaman I, Blakeley P, et al. GWAS for urinary sodium and potassium excretion highlights pathways shared with cardiovascular traits. Nat Commun. 2019 Aug 13;10(1):3653.

55. Finucane HK, Bulik-Sullivan B, Gusev A, Trynka G, Reshef Y, Loh P-R, et al. Partitioning heritability by functional annotation using genome-wide association summary statistics. Nat Genet. 2015 Nov;47(11):1228–35.

56. Sheng X, Guan Y, Ma Z, Wu J, Liu H, Qiu C, et al. Mapping the genetic architecture of human traits to cell types in the kidney identifies mechanisms of disease and potential treatments. Nat Genet. 2021 Sep;53(9):1322–33.

57. Poplin R, Ruano-Rubio V, DePristo MA, Fennell TJ, Carneiro MO, Auwera GAV der, et al. Scaling accurate genetic variant discovery to tens of thousands of samples [Internet]. 2018 Jul [cited 2022 Jan 6] p. 201178. Available from: https://www.biorxiv.org/content/10.1101/201178v3

58. Delaneau O, Zagury J-F, Robinson MR, Marchini JL, Dermitzakis ET. Accurate, scalable and integrative haplotype estimation. Nat Commun. 2019 Nov 28;10(1):5436.

59. Auton A, Abecasis GR, Altshuler DM, Durbin RM, Abecasis GR, Bentley DR, et al. A global reference for human genetic variation. Nature. 2015 Oct;526(7571):68–74.

60. van de Geijn B, McVicker G, Gilad Y, Pritchard JK. WASP: allele-specific software for robust molecular quantitative trait locus discovery. Nat Methods. 2015 Nov;12(11):1061–3.

61. Ma S, Zhang B, LaFave LM, Earl AS, Chiang Z, Hu Y, et al. Chromatin Potential Identified by Shared Single-Cell Profiling of RNA and Chromatin. Cell. 2020 Nov 12;183(4):1103-1116.e20.

62. Zimmerman KD, Espeland MA, Langefeld CD. A practical solution to pseudoreplication bias in single-cell studies. Nat Commun. 2021 Feb 2;12(1):738.

63. Fang R, Preissl S, Li Y, Hou X, Lucero J, Wang X, et al. Comprehensive analysis of single cell ATAC-seq data with SnapATAC. Nat Commun. 2021 Feb 26;12(1):1337.

64. Gutierrez-Arcelus M, Baglaenko Y, Arora J, Hannes S, Luo Y, Amariuta T, et al. Allele-specific expression changes dynamically during T cell activation in HLA and other autoimmune loci. Nat Genet. 2020 Mar;52(3):247–53.

65. Zhou Y, Luo Z, Liao C, Cao R, Hussain Z, Wang J, et al. MHC class II in renal tubules plays an essential role in renal fibrosis. Cell Mol Immunol. 2021 Nov;18(11):2530–40.

66. DeFronzo RA, Reeves WB, Awad AS. Pathophysiology of diabetic kidney disease: impact of SGLT2 inhibitors. Nat Rev Nephrol. 2021 May;17(5):319–34.

67. Kirita Y, Wu H, Uchimura K, Wilson PC, Humphreys BD. Cell profiling of mouse acute kidney injury reveals conserved cellular responses to injury. PNAS. 2020 Jul 7;117(27):15874–83.

68. Chang-Panesso M, Kadyrov FF, Lalli M, Wu H, Ikeda S, Kefaloyianni E, et al. FOXM1 drives proximal tubule proliferation during repair from acute ischemic kidney injury. J Clin Invest. 129(12):5501– 17.

69. Witchel SF, DeFranco DB. Mechanisms of disease: regulation of glucocorticoid and receptor levels--impact on the metabolic syndrome. Nat Clin Pract Endocrinol Metab. 2006 Nov;2(11):621–31.

70. Nissen RM, Yamamoto KR. The glucocorticoid receptor inhibits NFκB by interfering with serine-2 phosphorylation of the RNA polymerase II carboxy-terminal domain. Genes Dev. 2000 Sep 15;14(18):2314–29.

71. Sundahl N, Bridelance J, Libert C, De Bosscher K, Beck IM. Selective glucocorticoid receptor modulation: New directions with non-steroidal scaffolds. Pharmacol Ther. 2015 Aug;152:28–41.

72. De Bosscher K, Van Craenenbroeck K, Meijer OC, Haegeman G. Selective transrepression versus transactivation mechanisms by glucocorticoid receptor modulators in stress and immune systems. European Journal of Pharmacology. 2008 Apr 7;583(2):290–302.

73. Sasaki M, Sasako T, Kubota N, Sakurai Y, Takamoto I, Kubota T, et al. Dual Regulation of Gluconeogenesis by Insulin and Glucose in the Proximal Tubules of the Kidney. Diabetes. 2017 Sep;66(9):2339–50.

74. Tiwari S, Singh RS, Li L, Tsukerman S, Godbole M, Pandey G, et al. Deletion of the Insulin Receptor in the Proximal Tubule Promotes Hyperglycemia. JASN. 2013 Aug 1;24(8):1209–14.

75. Kamm DE, Fuisz RE, Goodman AD, Cahill GF. Acid-Base Alterations and Renal Gluconeogenesis: Effect of pH, Bicarbonate Concentration, and PCO2*. J Clin Invest. 1967 Jul;46(7):1172–7.

76. Inaba Y, Hashiuchi E, Watanabe H, Kimura K, Sato M, Kobayashi M, et al. Hepatic Gluconeogenic Response to Single and Long-Term SGLT2 Inhibition in Lean/Obese Male Hepatic G6pc-Reporter Mice. Endocrinology. 2019 Dec 1;160(12):2811–24.

77. Gronda E, Jessup M, Iacoviello M, Palazzuoli A, Napoli C. Glucose Metabolism in the Kidney: Neurohormonal Activation and Heart Failure Development. Journal of the American Heart Association. 2020 Dec 1;9(23):e018889.

78. Fan J, Hu J, Xue C, Zhang H, Susztak K, Reilly MP, et al. ASEP: Gene-based detection of allele-specific expression across individuals in a population by RNA sequencing. PLoS Genet. 2020 May;16(5):e1008786.

79. Thibodeau A, Eroglu A, McGinnis CS, Lawlor N, Nehar-Belaid D, Kursawe R, et al. AMULET: a novel read count-based method for effective multiplet detection from single nucleus ATAC-seq data. Genome Biology. 2021 Sep 1;22(1):252.

80. Corces MR, Trevino AE, Hamilton EG, Greenside PG, Sinnott-Armstrong NA, Vesuna S, et al. An improved ATAC-seq protocol reduces background and enables interrogation of frozen tissues. Nat Methods. 2017 Oct;14(10):959–62.

81. Bates D, Mächler M, Bolker B, Walker S. Fitting Linear Mixed-Effects Models Using lme4. Journal of Statistical Software. 2015 Oct 7;67:1–48.

