## Supplemental Material for "Multimodal single cell sequencing of human diabetic kidney disease implicates chromatin accessibility and genetic background in disease progression"

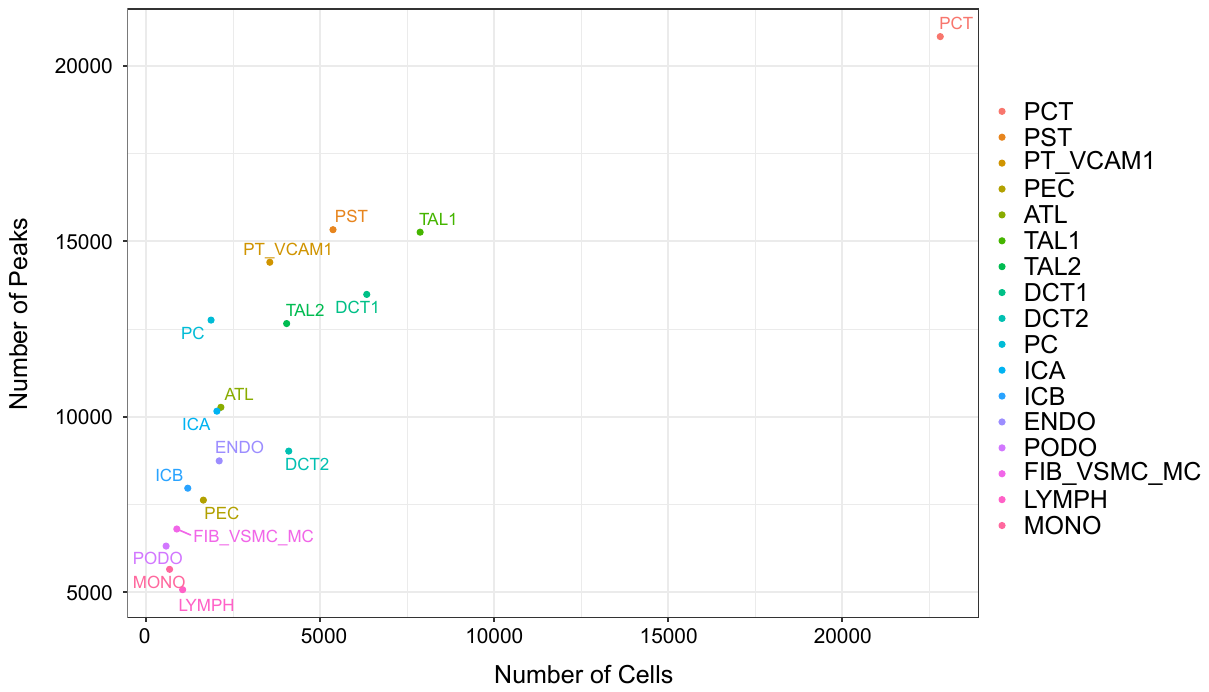

**Supplemental Figure 1. Number of MACS2 ATAC peaks called for each cell type.** Individual snATAC-seq cellranger-atac counts were aggregated for all thirteen libraries, preprocessed, and filtered with Signac. Cell-specific ATAC peaks were called separately for each cell type annotation using MACS2 via the Signac wrapper function. The number of cell-specific ATAC peaks meeting the adjusted p-value threshold (padj < 0.05) and the total number of cells passing quality control filters are visualized in a scatter plot.

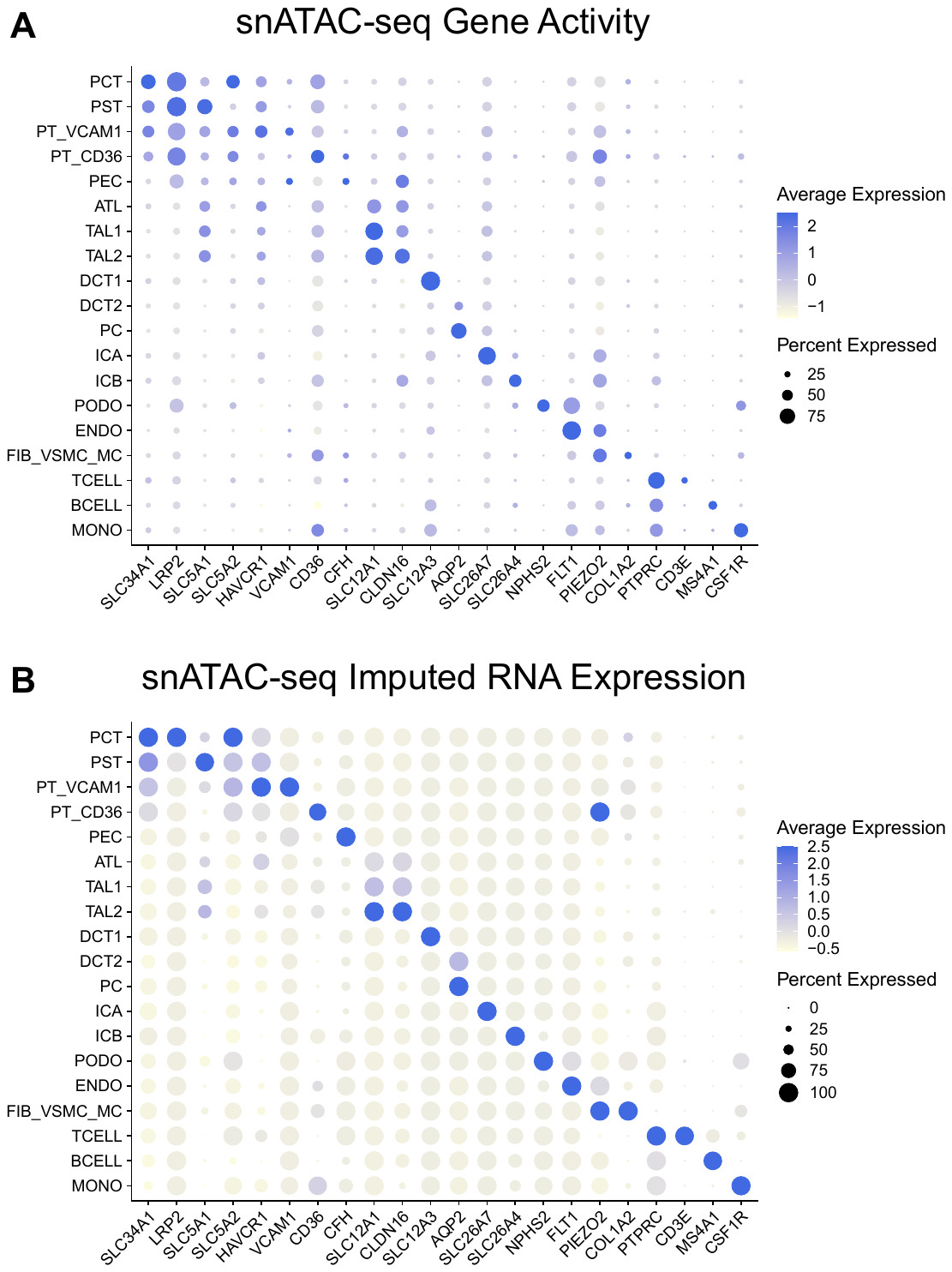

**Supplemental Figure 2. Lineage-specific markers in snATAC-seq. A)** Gene activity was computed for the aggregated and preprocessed snATAC-seq object for thirteen libraries using the gene body and promoter region of protein-coding genes and visualized with the Seurat DotPlot function. **B)** The aggregated snRNA-seq object was coembedded with the snATAC-seq object and estimated RNA expression values for the snATAC-seq object were computed using label transfer. Imputed RNA expression values are visualized with the Seurat DotPlot function.

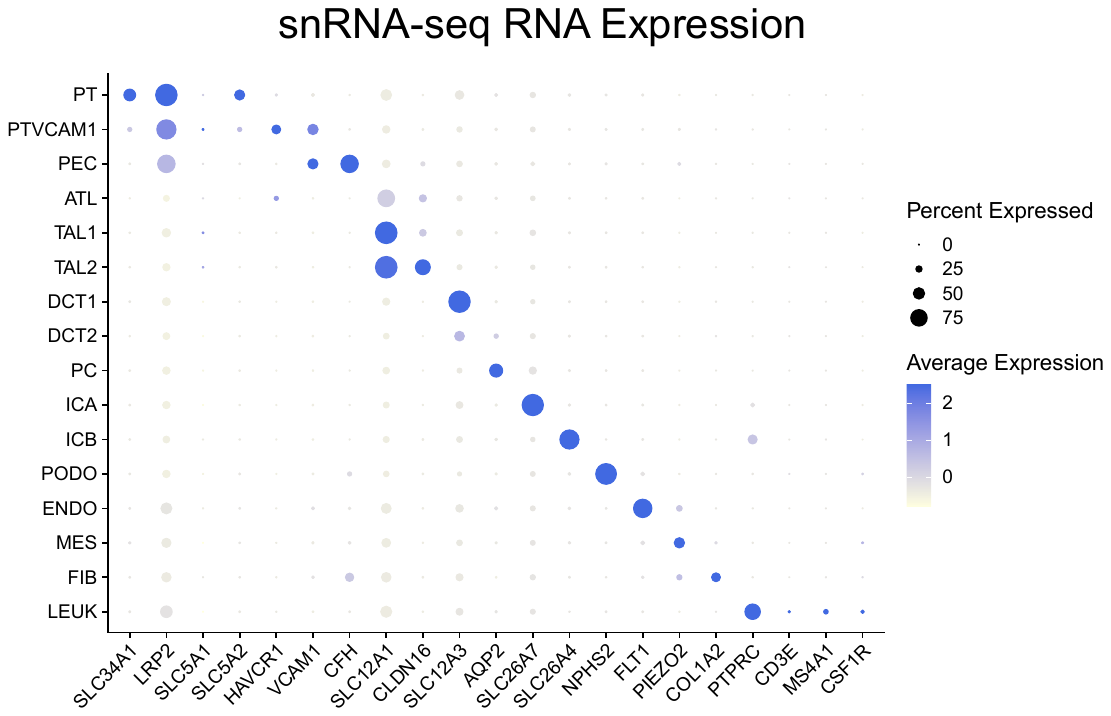

**Supplemental Figure 3. Lineage-specific markers by snRNA-seq.** Individual snRNA-seq cellranger libraries were aggregated, preprocessed, and filtered with Seurat. Lineage-specific markers for normalized RNA expression of individual cell types are visualized using the Seurat DotPlot function.
